# Astrocytic Signatures in Neuronal Activity: A Machine Learning-Based Identification Approach

**DOI:** 10.1101/2024.12.16.628802

**Authors:** João Pedro Pirola, Paige DeForest, Paulo R Protachevicz, Laura Fontenas, Ricardo F Ferreira, Rodrigo FO Pena

## Abstract

This study investigates the expanding role of astrocytes, the predominant glial cells, in brain function, focusing on whether and how their presence influences neuronal network activity. We focus on particular network activities identified as synchronous and asynchronous. Using computational modeling to generate synthetic data, we examine these network states and find that astrocytes significantly affect synaptic communication, mainly in synchronous states. We use different methods of extracting data from a network and compare which is best for identifying glial cells, with mean firing rate emerging with higher accuracy. To reach the aforementioned conclusions, we applied various machine learning techniques, including Decision Trees, Random Forests, Bagging, Gradient Boosting, and Feedforward Neural Networks, the latter outperforming other models. Our findings reveal that glial cells play a crucial role in modulating synaptic activity, especially in synchronous networks, highlighting potential avenues for their detection with machine learning models through experimental accessible measures.

## 1 Introduction

Glial cells are a prominent element that contribute to several brain functions (He and Sun, 2007; Parpura et al, 2012). These essential non-neuronal elements in the nervous system communicate with neurons through a complex network of interactions. They provide insulatory, structural, metabolic, and trophic support to neurons and are involved in regulating neurotransmitter levels and modulating synaptic transmission (Jessen, 2004; Jäkel and Dimou, 2017). Understanding the interactions between glial cells and neurons is crucial for gaining insights into how the brain maintains its function in various states, from health to disease (Linne et al, 2022; Adamczyk, 2023). This understanding can contribute to the development of computational models that simulate or identify these interactions, helping us decode the complex patterns of neural activity.

Among the diversity of glial cell subtypes, astrocytes are the most abundant in the brain (Jäkel and Dimou, 2017). Astrocytes have emerged as key regulators of neural network function, challenging the long-held neuron-centric view of brain activity. Traditionally regarded as passive support cells, astrocytes are now recognized as active participants in synaptic modulation, metabolic coupling, and the regulation of extracellular ionic and neurotransmitter homeostasis. By bridging vascular and neuronal networks, astrocytes contribute to dynamic processes such as synaptic plasticity, neurovascular coupling, and the orchestration of rhythmic network activity (Halassa and Haydon, 2010; Lines et al, 2020; Stackhouse and Mishra, 2021; Van Horn et al, 2021). They are also involved in the regulation and maintenance of the blood-brain barrier (Ben Achour and Pascual, 2012; Von Bartheld et al, 2016). However, despite these major findings, the complex interplay between astrocytes and neural networks remains poorly understood, partly due to the lack of integrative computational frameworks tailored to astrocytic mechanisms. Astrocytes communicate through calcium signaling rather than action potentials, creating a distinct mode of interaction with neural circuits. These calcium transients are influenced by local synaptic activity, vascular signals, and intrinsic astrocytic properties, allowing astrocytes to operate as central hubs of neural information processing (Straub and Nelson, 2007; Guerra-Gomes et al, 2018). Emerging evidence suggests that astrocytes do not simply modulate neuronal activity but also influence network stability, synaptic remodeling, and the synchronization of large-scale neural oscillations (Bazargani and Attwell, 2016; Poskanzer and Yuste, 2016; Perez-Catalan et al, 2021; Lawal et al, 2022; Kim and Chung, 2023).

Interestingly, there is also a growing body of evidence suggesting the involvement of astrocytes or other glial cells in the development and progression of neurodegenerative disorders in which memory and cognitive problems are prominent. In Alzheimer’s disease (AD), astrocytes undergo a range of changes, including alterations in gene expression, morphology, and function (Birch, 2014; Cai et al, 2017). Studies have indicated that astrocytes in AD may become less responsive to neuronal signaling, leading to the dysregulation of synaptic function and, eventually, neuronal death. Additionally, astrocytes in AD have been shown to exhibit impaired clearance of toxic proteins, such as amyloid-beta. These proteins can build up and cause plaques, which are a hallmark of the disease (Cai et al, 2017). At the same time, recent studies have emphasized heterogeneity among astrocytes within the central nervous system, with distinct subtypes exhibiting unique molecular signatures and functions (Bayraktar et al, 2015; Chen et al, 2020). Considering this diversity is crucial for understanding their specific roles in various brain regions and under different physiological and pathological conditions.

Clearly, glial cells are significantly implicated in several brain functions making identifying their presence among neurons an appealing problem (Dimou and Götz, 2014). To that end, modeling can be helpful. Nevertheless, the simulation of the complex interactions between glial cells and neurons is a challenging task that requires advanced computational approaches. For example, the work of De Pitt`a and Brunel has shown that astrocyte-mediated gliotransmission can be an important player in modulating working memory (WM) (De Pitt`a and Brunel, 2022). Sufficiently strong stimulation in subnetworks composed of excitatory and inhibitory cells with glial feedback creates multiple attractors that act independently. At the level of a single subpopulation, there were multiple equilibrium points reported (Wei and Shuai, 2011; Wang et al, 2012). Overall, the role of glial cells in multiple brain patterns has been observed through synchronization (Kanner et al, 2018; Handy and Borisyuk, 2023), spatial correlations (Handy and Borisyuk, 2023), synaptic plasticity (Gómez-Gonzalo et al, 2015; Sherwood et al, 2017), synaptic pruning (Stevens et al, 2007; Paoli-celli et al, 2011), and modulation of neuronal signals (Jackson, 2011; Newman, 2015; Purushotham and Buskila, 2023). However, most of these works consider synchronous states and do not address network pattern changes and how glial cell feedback affects them.

In this paper, we are interested in identifying astrocytes using neuronal measures in both synchronous and asynchronous states. By utilizing models that account for both synchronous and asynchronous neural states, we can better understand the influence of glial cells on different brain network dynamics. From the experiments that were conducted, we separated four cases: two synchronous and two asynchronous. These cases differ in terms of external stimulation or conductance level and serve as the basis for the creation of an artificial dataset with many samples. Since the identification of the presence of glial cells could be dependent on the recorded signal, we investigated different experimental collection methods, the generated data included two of them: one that collects mean firing rate and the other where voltage count is obtained. Our goal was to identify the presence of glial cells in synaptic transmission using different machine-learning methods, which do not require strong assumptions about the data.

To achieve this goal we systematically used several machine learning methods including Feedforward Neural Networks (FNNs) (Rosenblatt, 1958), Decision Trees (Breiman et al, 1984a), Bagging (Breiman, 1996), Random Forests (Breiman, 2001), and Gradient Boosting (Friedman, 2001). These models were selected for their effectiveness in handling multicollinearity and suitability for classification tasks. Our results show that FNNs performed better for asynchronous states using mean firing rate, whereas when voltage count is used as the data collection method, FNNs performed similarly to other learning algorithms. We optimized algorithm performances using Bayesian optimization (Wu et al, 2019) and cross-validation, ensuring robust hyperparameter selection and reliable findings.

Our work is organized as follows: In Subsection 2.1, we discuss the neuron and astrocyte models used in our study. Subsection 2.2 covers the measures employed to analyze network dynamics, particularly regarding synchronous and asynchronous states, voltage count, and firing rate. In Subsection 2.3, we present the selected machine learning classifiers. Subsection 2.4 outlines the performance measures used to evaluate the classifiers. Section 3 provides an in-depth description of our results, beginning with the essential dynamics for characterizing network activity with and without glial cells, with key metrics such as firing rate and coefficient of variation (CV), analyzing synchronous and asynchronous states through specific parameter combinations. We investigate how astrocytes generate a broader distribution of spike train coherence, focusing on individual cell communication. Lastly, in subsection 3.5, we detail the machine learning classifier results. Section 4 presents a comprehensive discussion and final remarks.

## 2 Methods

### 2.1 Computational models

#### 2.1.1 Neuron Model

The dynamics of each single neuron is described by the conductance-based leaky integrate-and-fire (LIF) model, which captures essential biophysical processes underlying neuronal activity (Burkitt, 2006). The membrane potential *V* (*t*) of a neuron at time index *t* evolves according to the following differential equation

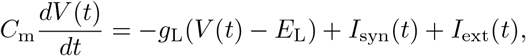

where *C*_m_ represents the membrane capacitance, defining the neuron’s ability to store charge, *g*_L_ is the leak conductance, controlling the passive flow of ions through the membrane, and *E*_L_ is the leak reversal potential, setting the resting membrane potential (Gerstner et al, 2014). The chemical synaptic current, *I*_syn_ (*t*), results from the input of other neurons via synapses, while *I*_ext_ (*t*) represents external inputs such as sensory stimuli or experimentally applied currents (Protachevicz et al, 2020).

In this model, the neuron fires an action potential when the membrane potential exceeds a threshold *V*_th_ (Destexhe, 1997). Upon reaching this threshold, a rapid depolarization occurs, followed by a reset mechanism that returns the potential to a fixed value, *V*_reset_ (Cessac and Viéville, 2008). To effectively model the refractory period in IF models, we consider that the membrane potential stays the rest for *τ*_r_ = 5 ms. The LIF model captures the essence of neuron behavior in a computationally efficient manner, making it widely used in large-scale network simulations (Sanaullah et al, 2023). The neuronal parameters considered in this work are given in Table S2.

#### 2.1.2 Astrocyte Model

Astrocytes play a crucial role in modulating synaptic transmission, primarily through the regulation of neurotransmitter release and uptake (Covelo and Araque, 2018; Purushotham and Buskila, 2023). The model of astrocytic activity focuses on intracellular calcium (Ca^2+^) dynamics, which are key to the communication between astrocytes and neurons (Fellin, 2009). The concentration of calcium within an astrocyte, [Ca^2+^], is described by a differential equation that responds to synaptic activity

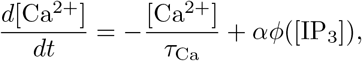

where *τ*_Ca_ represents the calcium decay time constant, dictating how quickly calcium levels return to baseline after an influx. The rate of calcium influx is governed by *α*, and *ϕ* ([IP_3_]) is a nonlinear function that determines the calcium release based on inositol trisphosphate ([IP_3_]) levels. [IP_3_] is generated in response to synaptic activity and mediates the release of calcium from intracellular stores, creating a feedback mechanism where astrocytes regulate synaptic behavior (Liu et al, 2021; Sherwood et al, 2021).

This model captures how astrocytes influence synaptic transmission through calcium-dependent processes, modulating the release of gliotransmitters that can enhance or inhibit neuronal firing. As calcium levels rise, they can alter synaptic strength, thus playing a role in neuroplasticity. For more intricate details regarding astrocyte-neuron interactions, refer to (Li and Rinzel, 1994; Perea and Araque, 2007; Panatier et al, 2011; Stimberg et al, 2019; De Pitt`a and Brunel, 2022). In the following sections, we will use the term glial cells or astrocytes interchangeably.

#### 2.1.3 Synaptic Interaction

The synaptic current *I*_syn_(*t*) received by a neuron at time *t* results from the interaction between presynaptic and postsynaptic neurons and is modeled as

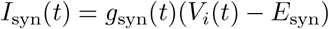

where *g*_syn_ (*t*) represents the synaptic conductance (Shimoura et al, 2021) and which is dynamically modulated by astrocytic activity. Synaptic conductance determines the strength of the synaptic connection at time *t*, and *E*_syn_ is the synaptic reversal potential, representing the voltage at which the synaptic current reverses direction.

Synaptic conductance *g*_syn_ (*t*) increases by a fixed amount in response to presynaptic spikes, simulating the release of neurotransmitters into the synaptic cleft. After each increase, *g*_syn_ (*t*) decays exponentially with a time constant that reflects the duration of synaptic influence. This process models the transient nature of synaptic transmission and how it is modulated by surrounding astrocytes. Immediately after the spike of the pre-synaptic neuron, *g*_syn_ is added to its value of 0.05 nS (excitatory) and 1.0 nS (inhibitory). The decay constant defines how quickly the synaptic strength diminishes over time, ensuring that the influence of each synaptic event is temporally limited.

Incorporating astrocyte-modulated synaptic conductance adds a layer of complexity, reflecting the bidirectional communication between neurons and astrocytes, where astrocytic calcium dynamics can enhance or suppress synaptic efficacy depending on the physiological state of the network (Sanz-Gálvez et al, 2024). This astrocyte-neuron interaction is critical for understanding mechanisms underlying learning and memory, particularly synaptic plasticity (Kol et al, 2020).

#### 2.1.4 Network

In this study, we modeled a network composed of excitatory and inhibitory neurons, coupled with astrocytes to investigate their influence on neuronal activity. The network was designed to capture key aspects of biological neural dynamics, incorporating both neuron-to-neuron and neuron-to-astrocyte interactions. We assume 1000 excitatory neurons and 250 inhibitory neurons, as well as 2000 astrocytes, unless stated otherwise. These cells are randomly connected where the excitatory cells connect to all other neurons with 0.05 probability, the inhibitory cells with 0.2, and astrocytes connect to at least one excitatory cell (De Pitt`a and Brunel, 2022).

### 2.2 Measures

In this subsection, we introduce the measures used to assess the properties which characterize the neuronal network dynamics. We consider two primary metrics for classification: one derived from spiking activity and another from voltage activity.

To monitor the spiking activity, we selected the mean firing rate, a widely used metric in both experimental and computational neuroscience. The mean firing rate provides a straightforward yet effective means of quantifying the average spiking activity of neurons over a defined time window. This measure aligns with real-world brain data collection methodologies, where firing rates are analyzed to understand the overall activity levels of neural networks. By averaging spike counts across all neurons, the mean firing rate serves as a robust indicator of network activity, making it well-suited for assessing large-scale neural systems.

For voltage activity, we used the voltage count, or distribution, across all neurons in the network. This method resembles techniques used to capture the potential of specific brain regions, offering insights into the likelihood of neurons maintaining particular voltage levels. To provide a comprehensive characterization of the network, we also included the coefficient of variation (CV) of interspike intervals and pairwise coherence between neurons.

These measures can offer additional insights into the variability and synchrony of neuronal firing, respectively. Further details on each of these methods are provided in the following.

#### 2.2.1 Voltage distribution

We calculated the averaged voltage distribution by performing a voltage count across all neurons, similar to constructing a histogram of voltage values. This approach provides the count, or probability, of observing a neuron at a particular voltage level, averaged across the network. By binning voltage values and counting occurrences in each bin, we obtained a distribution representing the likelihood of finding a neuron at a specific voltage.

#### 2.2.2 Firing rate

We calculated the mean firing rate (Hz) of all neurons in the network by

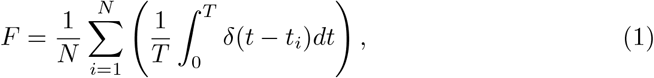

where *t*_*i*_ is the time of the *i*-th spike, *N* is the number of neurons in the network, and *T* = 10s is the time window considered for analyses (Tomov et al, 2016).

#### 2.2.3 Coefficient of variation (CV)

The CV provides information about the irregularity of neuronal activities which can be applied from single to neuronal networks. To obtain the CV, we first calculated the inter-spike intervals (ISI) where the *i*th interval is defined as the difference between two consecutive spike times *t*_*i*+1_ and *t*_*i*_, namely ISI_i_ = *t*_*i*+1_ − *t*_*i*_ *>* 0 (Protachevicz et al, 2022). From the ISI series, the first interval is referred to as ISI_1_ followed by the subsequent intervals, namely ISI_2_, ISI_3_, …, and ISI_n_. The ratio between the standard deviation and the mean (indicated by ⟨ ·⟩) gives rise to the coefficient of variation (CV_*i*_) for the *ith* neuron (Lengler and Steger, 2017):

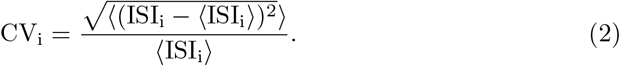

Finally, the average of CV_*i*_ of all *N* neurons is given by

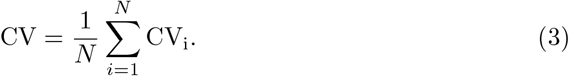

The range of CVs can vary, and it is assumed to be close to 1 for a Poisson process. Values lower than 1 will indicate a more periodic activity.

#### 2.2.4 Coherence

To characterize the frequency-dependent communication of neurons, we consider the coherence measurement which is associated with the level of similarity of two signals at some frequency. By defining the spike train *x*(*t*) as the sum of *δ* functions at spike times *t*_*i*_ (Borges et al, 2020)

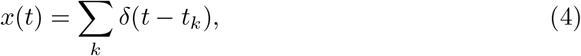

we could obtain the power spectrum *S*_*xx*_ of *x*(*t*) with

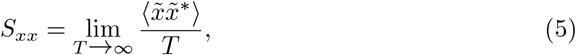

where we used the Fourier transformation of *x*(*t*) given by 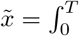 *dt* exp(*iωt*)*x*(*t*) and its correspondent conjugate 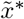. For spike trains selected from different neurons, *x*(*t*) and *y*(*t*), we have the corresponding cross-spectra *S*_*xy*_.

Then, the frequency-dependent correlations between *x*(*t*) and *y*(*t*) could be determined by the coherence function:

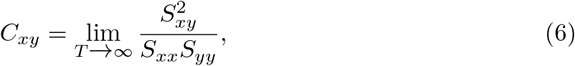

where, for convenience, we present the averaged value over all the frequency content for each pair of neurons (Pena et al, 2018).

### 2.3 Classifiers

In this section, we briefly describe the machine learning methods used to identify the presence of glial cells in synaptic transmission. We chose the following models—Feedforward Neural Networks (FNNs), Decision Trees, Bagging, Random Forests, and Gradient Boosting—because we observed strong multicollinearity among the variables, and these models do not make strong assumptions about independence between features. We also refer the reader to Appendix A where we explain in depth our choice and initialization of parameters and hyperparameters for each of these classifiers.

#### 2.3.1 Decision Trees

A Decision Tree (DTree) (Breiman et al, 1984a) is a supervised machine learning algorithm used for both classification and regression tasks, which splits data into subsets based on the value of input features. It constructs a tree-like structure where each internal node represents a decision (based on a feature) and each leaf node represents an outcome (either a class label or a regression value). In the case of classification, the algorithm selects the feature that best separates the data at each node using criteria like Gini Impurity (Gini, 1912) or Information Gain (Shannon, 1948). The tree grows recursively by splitting the data until a stopping condition is met.

##### Pseudocode

~~~
1. Input: Training data D, stopping criteria (max depth, min samples, etc.)
2. Procedure:
 a. Check stopping criteria. If met, stop and return a leaf node.
 b. For each feature, calculate the splitting criterion (e.g., Gini).
 c. Select the best feature and its threshold that maximizes the criterion.
 d. Split the data D into two subsets:
   D_left = data where feature <= threshold
   D_right = data where feature > threshold
 e. Create a decision node with the selected feature and threshold.
 f. Recursively repeat steps (a)-(e) for D_left and D_right to build tree.
3. Output: Decision tree.
~~~

#### 2.3.2 Bagging

Bagging (Bootstrap Aggregating), first introduced by Breiman (1996) is an ensemble learning technique that improves the performance of models like Decision Trees by reducing variance. It works by generating multiple subsets of the training data through random sampling with replacement (bootstrap sampling). A Decision Tree model is trained on each subset, and the final prediction is made by aggregating the results from all models, typically using majority voting for classification or averaging for regression. By combining the predictions from multiple trees, Bagging increases stability and accuracy, particularly for models prone to overfitting.

##### Pseudocode

~~~
1. Input: Training data D, number of models T, base learner
     (e.g., Decision Tree)
2. Procedure:
 a. For each model i from 1 to T:
  i. Generate a bootstrap sample D_i by randomly sampling with replacement from D.
  ii. Train the base learner (e.g., a Decision Tree) on the bootstrap sample D_i.
 b. For classification: Aggregate predictions from each model
    by majority voting.
    For regression: Aggregate predictions from each model
    by averaging.
3. Output: Final prediction based on aggregated results.
~~~

#### 2.3.3 Random Forest

Random Forest (RF) (Breiman, 2001) is an ensemble learning method used for classification and regression. It builds multiple decision trees during training and aggregates their predictions. Each tree is trained on a random subset of the data using bootstrap sampling and a random subset of features. This randomness helps reduce overfitting and variance, making the model more robust.

##### Pseudocode

~~~
1. Input: Training data D, number of trees T, number of features F.
2. Procedure:
 a. For each tree t in range(T):
  i. Sample a subset of data D_t from D (with replacement).
  ii. Select a random subset of features F_t from F.
  iii. Build a decision tree on D_t using F_t.
 b. Aggregate predictions from all T trees (e.g., majority
    vote for classification).
3. Output: Random Forest model.
~~~

#### 2.3.4 Gradient Boosting

Gradient Boosting (GBoost) (Friedman, 2001) is another ensemble technique where models are built sequentially. Each new model tries to correct the errors of the previous one by optimizing a loss function. Unlike Random Forest, where trees are independent, Gradient Boosting builds trees that are dependent, with each one learning from the residual errors of its predecessors.

##### Pseudocode

~~~
1. Input: Training data D, number of trees T, learning rate (h).
2. Procedure:
  a. Initialize the model f(x) with a simple prediction (e.g., mean).
  b. For each tree t in range(T):
   i. Calculate residual errors: r_t = y - f(x) (for each instance).
   ii. Fit a decision tree tree(x) to the residuals r_t.
   iii. Update the model: f(x) = f(x) + h * tree(x).
  c. Repeat until T trees are built or stopping criteria are met.
3. Output: Gradient Boosting model.
~~~

#### 2.3.5 Feedforward Neural Networks

Feedforward Neural Networks (FNNs) (Rosenblatt, 1958) are a class of Artificial Neural Networks (McCulloch and Pitts, 1943) where connections between the nodes do not form cycles. These networks consist of multiple layers: an input layer, one or more hidden layers, and an output layer. Each node (or neuron) in a layer is connected to every node in the next layer via weights. The network learns by adjusting these weights based on the error between predicted and actual outcomes.

We consider a network with an input vector (*X*_1_, …, *X*_*p*_), where each *X*_*i*_ is a predictor, *i* = 1, 2, …, *p*. These inputs include information on the presence or absence of astrocytes. The nodes of the first hidden layer are represented by the vector 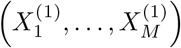, where each variable *X*^(1)^ is a function of a linear combination of the input vector elements. Specifically,

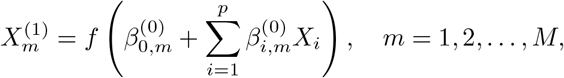

where *f* is a non-linear activation function, 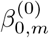 is the bias term, and 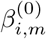 are weights connecting input nodes to the first hidden layer.

Once the values of 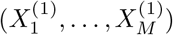 are computed, propagation continues sequentially through the network. For a multi-layer neural network with *H* hidden layers, each layer *h* has *d*_*h*_ nodes. The recursive formula for the hidden layer nodes follows the same pattern, involving a linear combination of the previous layer’s outputs followed by the activation function.

The output layer, which predicts *K* classes, uses similar equations. For each class *k* = 1, 2, …, *K*, the output node is defined as a linear combination of the final hidden layer outputs. In our case, where *K* = 2, the output is converted into a probability for binary classification.

To estimate the parameters of the FNN model, we employed different optimization algorithms. These techniques adjust the weights and biases to minimize a chosen loss function, which in our case was binary cross-entropy, a standard loss function for binary classification tasks (Ian Goodfellow, 2016).

##### Pseudocode

~~~
1. Input: Training data D, weights W, biases b, learning rate h and
       number of epochs n.
2. Procedure:
  a. Initialize weights W and biases b randomly.
  b. For each epoch (iteration):
   i. Forward pass: For each layer l, compute:
     z_l = W_l * a_(l-1) + b_l (weighted sum)
     a_l = activation(z_l) (non-linear transformation)
   ii. Compute the error E (e.g., cross-entropy for classification).
   iii. Backward pass: Propagate the error backward to compute gradients:
      dE/dW and dE/db using the chain rule.
   iv. Update weights and biases:
     W = W - h * dE/dW
     b = b - h * dE/db
3. Output: Trained neural network.
~~~

### 2.4 Performance measures

To quantify the model’s performance, we use validation data to compute the true positives (TP), true negatives (TN), false positives (FP), and false negatives (FN). These values form the confusion matrix, which is structured as follows.

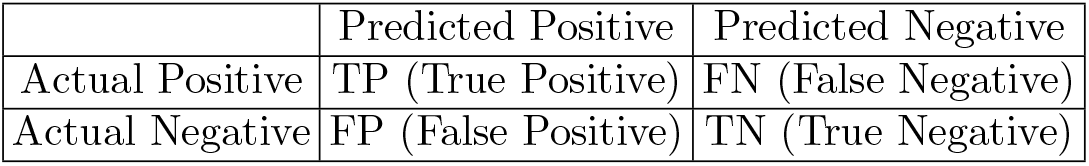

|  | Predicted Positive | Predicted Negative |
| --- | --- | --- |
| Actual Positive | TP (True Positive) | FN (False Negative) |
| Actual Negative | FP (False Positive) | TN (True Negative) |

From the confusion matrix, we can calculate several standard machine learning metrics that serve to characterize the model’s performance. These include:

#### Sensitivity (Recall)

Sensitivity (SN), also known as Recall, measures the model’s ability to correctly identify positive cases among all truly positive cases. The formula for Sensitivity is given by

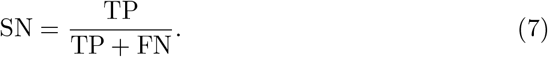

#### Specificity

Specificity (SP) measures the proportion of negative cases that are correctly identified by the model relative to the total number of truly negative cases. The formula for Specificity is given by

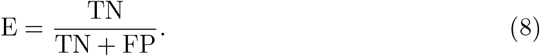

#### Accuracy

Accuracy (ACC) measures the proportion of correct predictions made by the model relative to the total number of samples. The formula for Accuracy is given by:

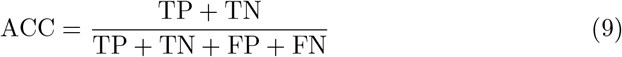

#### Positive Predictive Value (Precision)

Positive Predictive Value (PPV), also known as Precision, measures the proportion of correctly predicted positive cases relative to the total number of cases predicted as positive by the model. The formula for PPV is given by

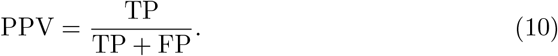

#### Negative Predictive Value (NPV)

Negative Predictive Value (NPV) measures the proportion of correctly predicted negative cases relative to the total number of cases predicted as negative by the model. The formula for NPV is given by

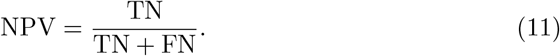

#### F1-Score

The F1-Score is the harmonic mean of Precision and Sensitivity (Recall). The formula for F1-Score is given by

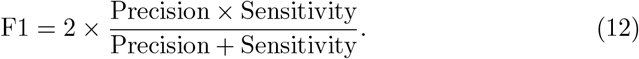

The F1-Score balances the model’s ability to avoid false positives and false negatives. A higher F1-Score indicates better performance, as it balances sensitivity and precision.

## 3 Results

### 3.1 Computational simulations reveal that astrocytic signatures in neuronal activity are of difficult identification

We begin by presenting key dynamics used to characterize network activity in the presence and absence of glial cells. Figure 7 illustrates the parameter spaces of firing rate and the average coefficient of variation (CV) across various input currents and excitatory weights for networks with and without glial cells. Our analysis centers on the firing rate and CV as primary indicators of network behavior.

**Fig 1.**
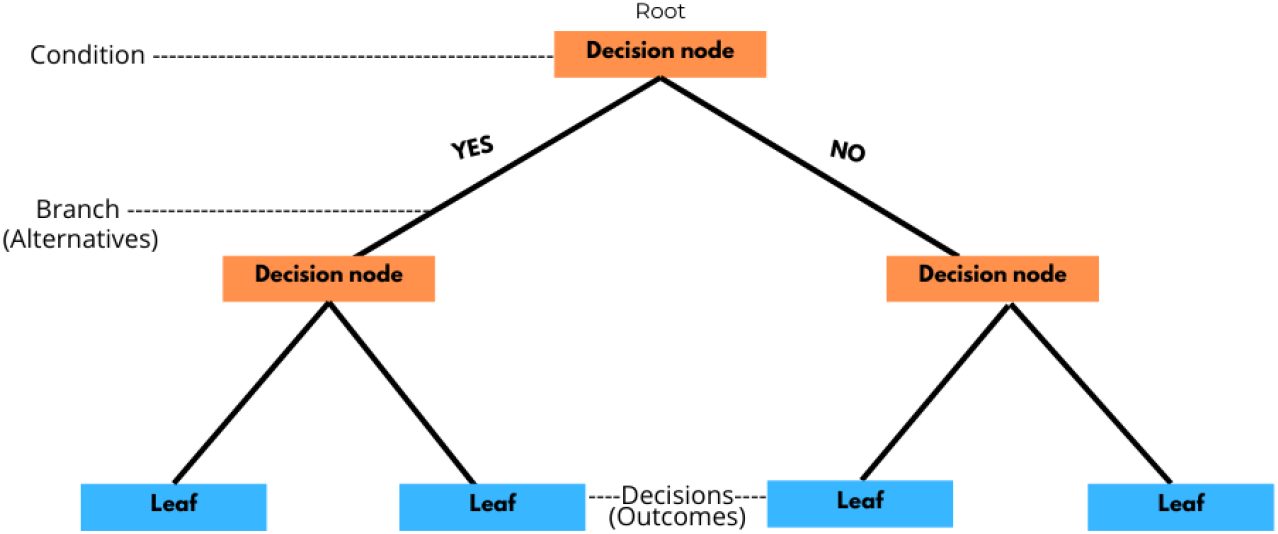
Decision Tree Structure Illustration. This figure shows the structure of a decision tree. The root node at the top represents the initial condition, leading to branches based on alternative outcomes. Each decision node (orange) represents a condition that splits the data based on specific criteria, leading to further branches. The branches represent possible alternatives, labeled here as “YES” and “NO.” The endpoints of each branch are the leaf nodes (blue), which provide the final decisions or outcomes. The decision tree model uses these nodes and branches to classify or predict outcomes based on input conditions.

**Fig 2.**
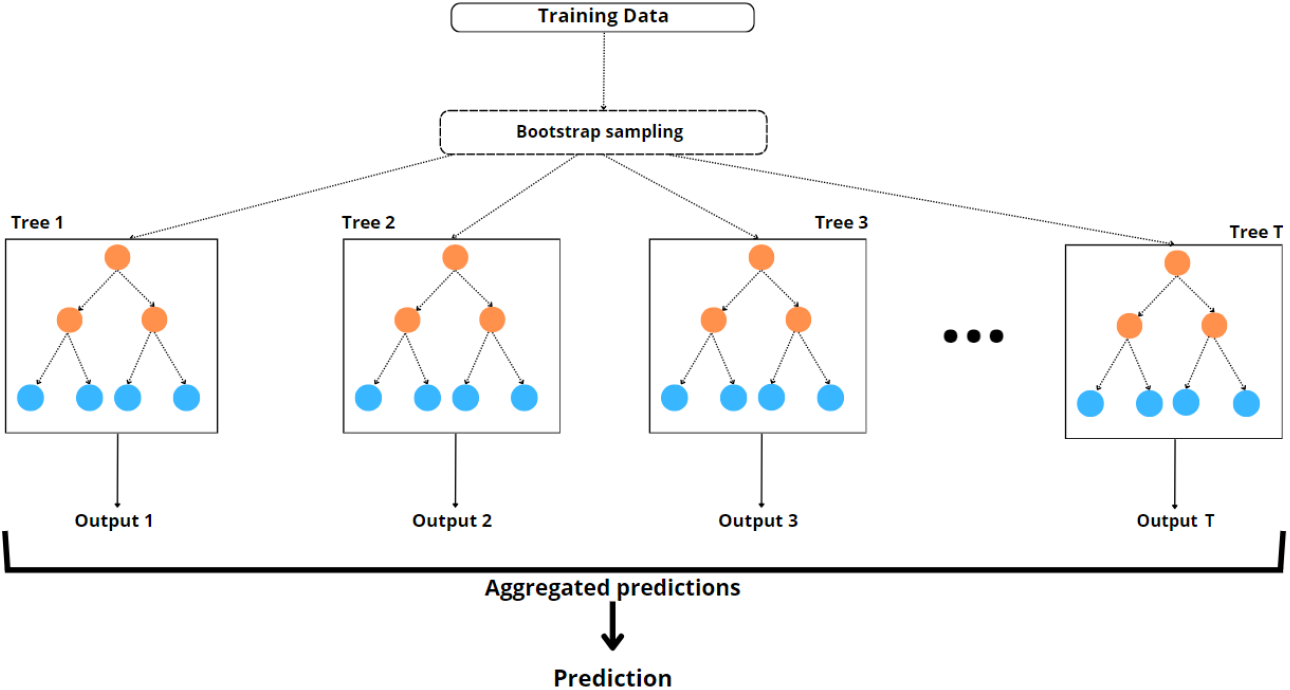
Bagging Algorithm Illustration. This figure shows the Bagging (Bootstrap Aggregating) process, where multiple decision trees are trained independently on different bootstrap samples of the training data. Bootstrap sampling is used to create unique datasets for each tree, allowing each tree to learn from a slightly different perspective. The orange nodes represent decision splits based on selected features, while the blue nodes are leaf nodes with the final predictions for each tree. Each tree produces an output (Output 1, Output 2, etc.), and these outputs are aggregated to generate the overall prediction.

**Fig 3.**
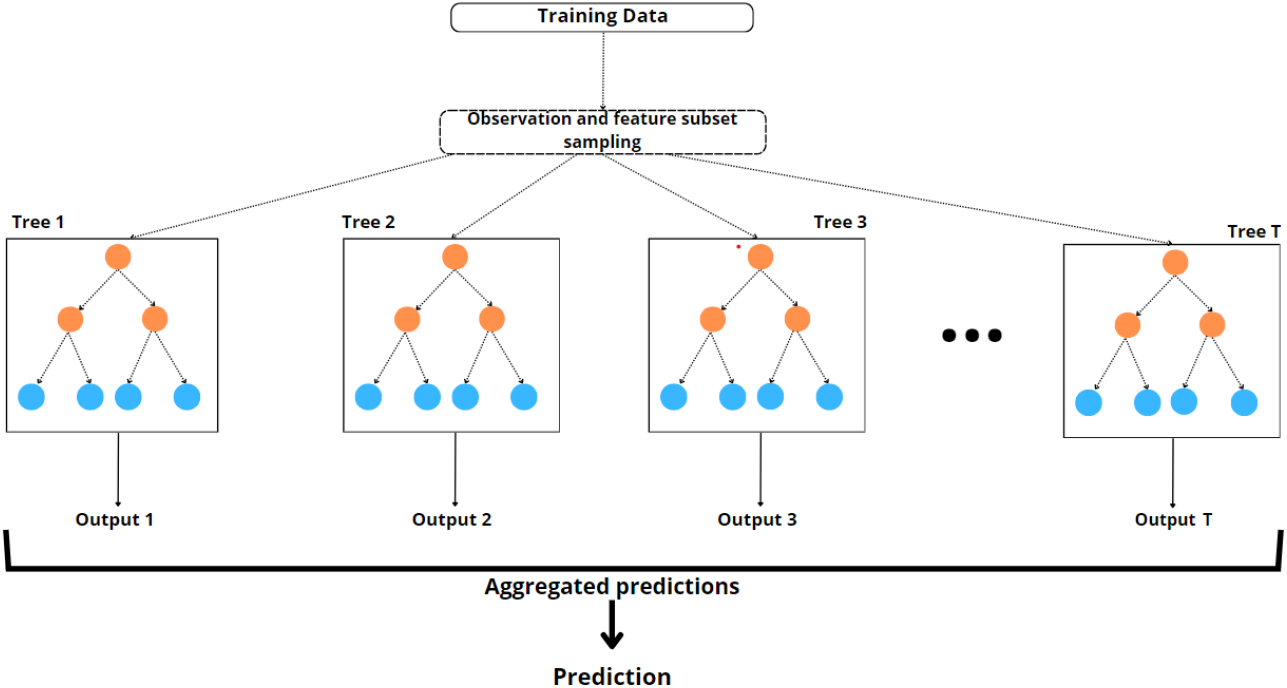
Random Forest Algorithm Illustration. This figure shows a Random Forest model, where multiple decision trees are trained in parallel on subsets of the training data. Each tree is built using a unique combination of observation and feature subset sampling. The orange nodes represent splits in the trees based on selected features, while the blue nodes represent leaf nodes, which contain the final outputs for each tree. The predictions from each tree (Output 1, Output 2, etc.) are aggregated to produce the overall model prediction.

**Fig 4.**
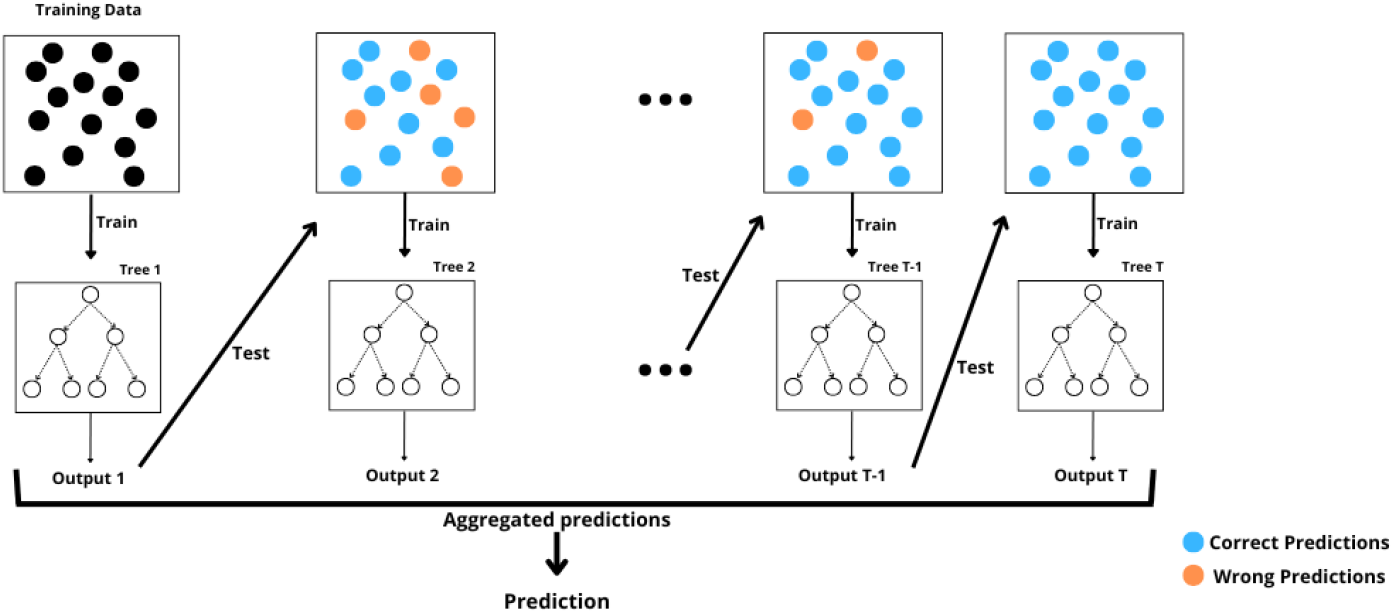
Gradient Boosting Algorithm Illustration: The figure shows a sequence of decision trees being trained iteratively. Each tree is trained on data with previous predictions, with orange points representing incorrect predictions and blue points representing correct predictions. The output from each tree is combined (aggregated predictions) to form the final prediction. Each subsequent tree aims to correct errors from the previous trees, enhancing model accuracy through boosting.

**Fig 5.**
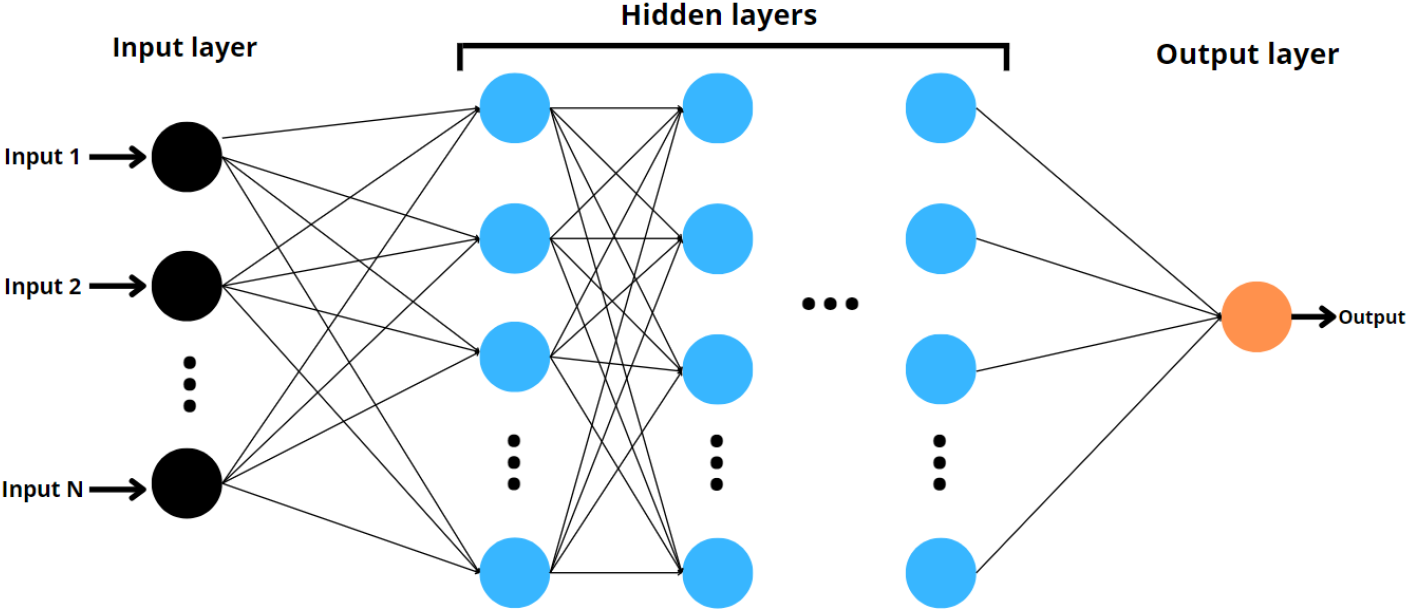
Feedforward Neural Network architecture. The figure shows an input layer as black circles, hidden layers as blue circles, and output layer as an orange circle. The circles are the artificial neurons and the arrows represent the connections between the neurons.

As seen in the figure, the presence of glial cells does not substantially alter the mean firing rate or CV across these parameter sets, with only slight variations observed in the CV under conditions of high input and low excitatory conductance. This subtlety highlights the difficulty of detecting glial cells using common network metrics, as the changes they induce are mild and easily overlooked. This reinforces the need for more sophisticated approaches to reliably identify glial influence within neuronal networks.

To understand the impact of astrocytes on network dynamics, we focus on four specific parameter combinations. These combinations are illustrated in Fig. 7. We selected cases of synchronous and asynchronous activity (top and bottom panels of Fig. 6). The time-dependent firing rate (black traces) remains steady after initial conditions stabilize in asynchronous cases, while it fluctuates in synchronous cases. When astrocytes are present, individual cell dynamics are depicted with green curves. For oscillatory (synchronous) cases, the oscillations become more pronounced when astrocytes are activated mid-simulation, suggesting a stronger astrocytic influence on oscillatory behavior.

**Fig 6.**
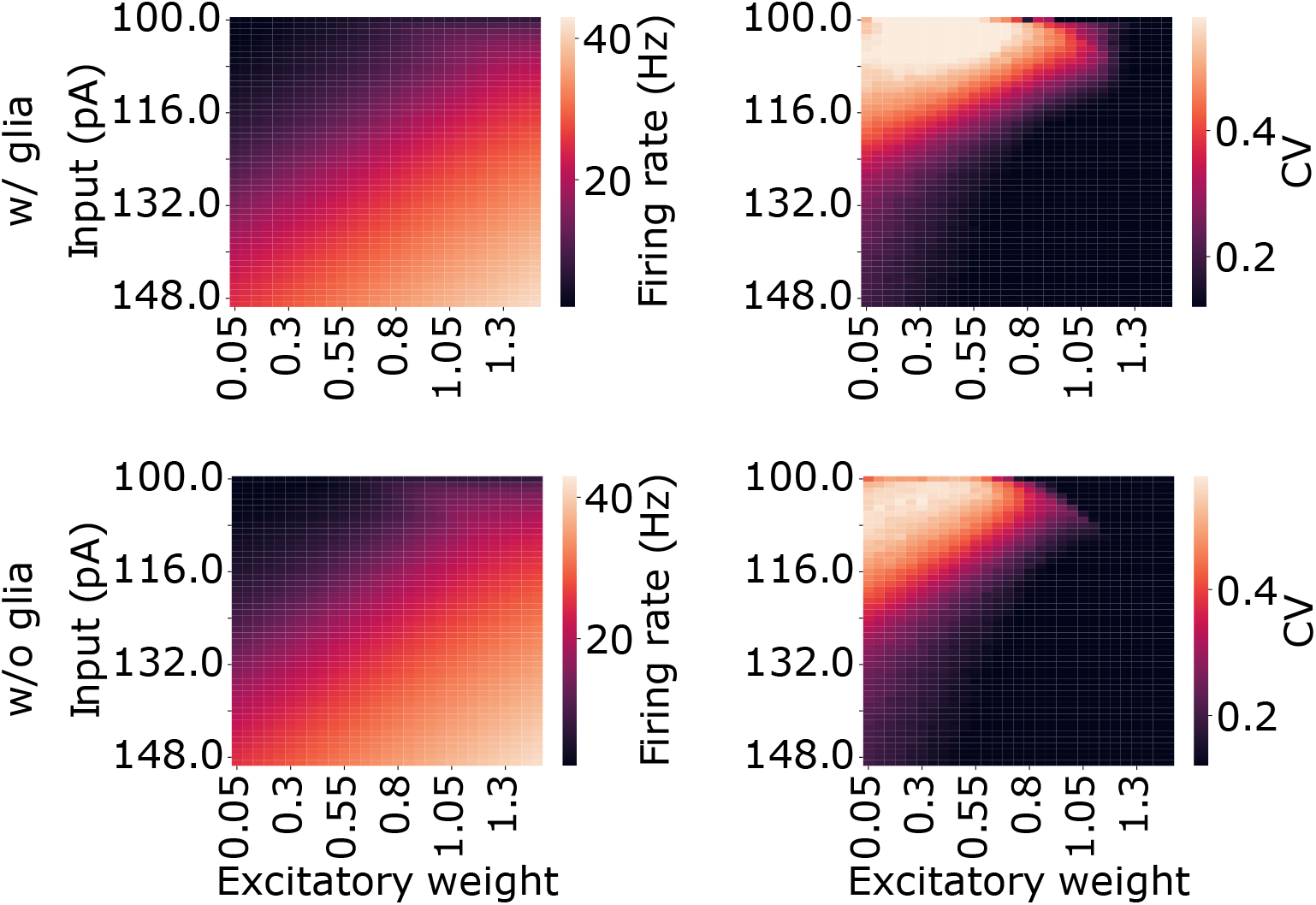
Parameter spaces for a random network with glial cells (top panels) and without glial cells (bottom panels), showing the effects of varying external stimulation (*I*_ext_) and excitatory weight (*g*_exc_). The colored diagrams represent both the network firing rate and coefficient of variation (CV).

**Fig 7.**
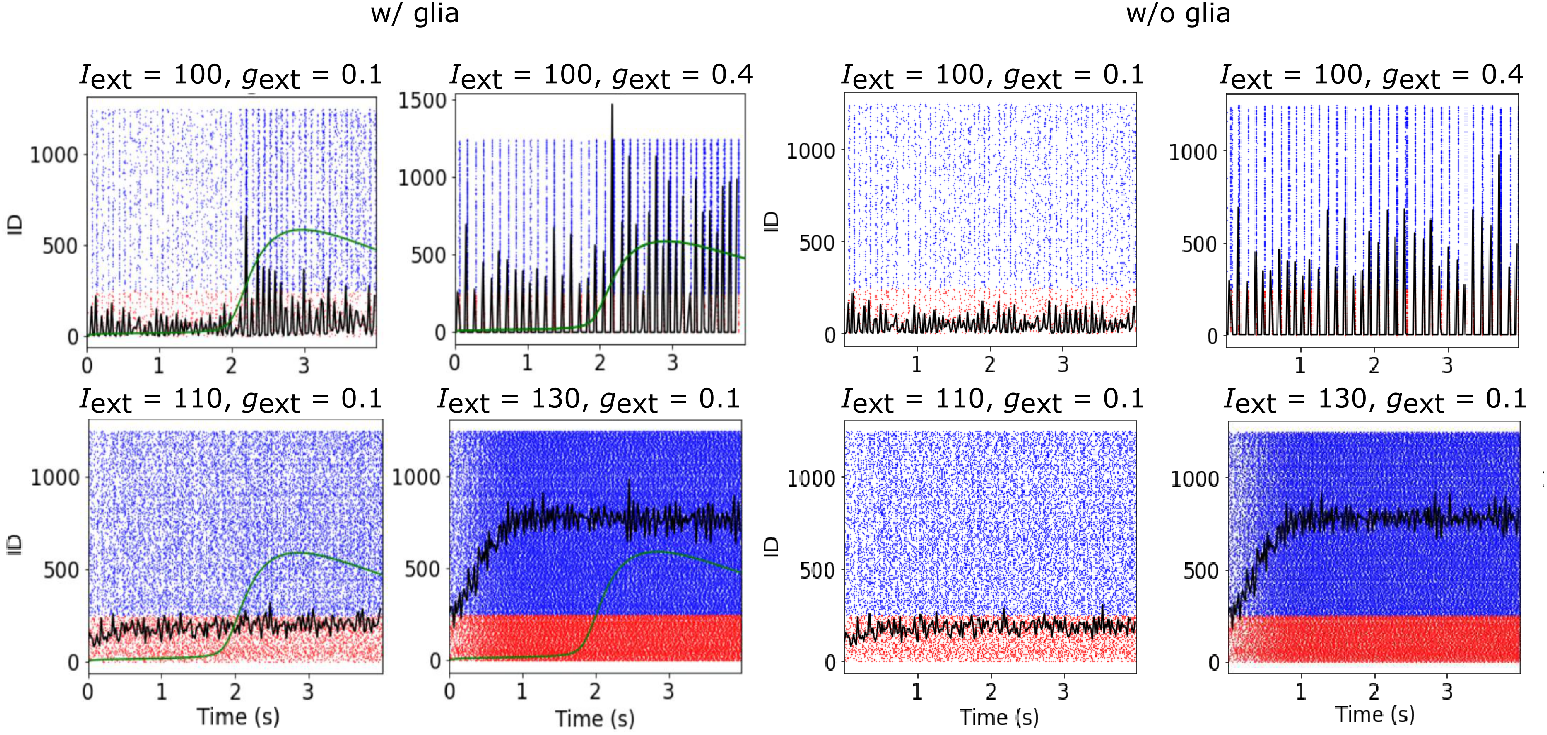
Selected exemplary network simulations from the parameter space in Fig. 6 (see values atop). Raster plots show both excitatory (blue) and inhibitory (red) populations down-sampled. Black curves: time-dependent network firing rate. Green curves: one example of calcium activation for one astrocyte randomly selected in the network. The top cases are considered synchronous whereas the bottom are asynchronous.

These patterns were further analyzed through the firing rate power spectrum, which confirmed that simulations with parameters (*I*_ext_ = 100 pA, *g*_exc_ = 0.1) and (*I*_ext_ = 100 pA, *g*_exc_ = 0.4) exhibit a peak in the spectrum, indicating synchronous behavior. In contrast, the cases with (*I*_ext_ = 110 pA, *g*_exc_ = 0.1) and (*I*_ext_ = 130 pA, *g*_exc_ = 0.1) lack this peak, characterizing an asynchronous regime. We will, therefore, refer to the first pair as synchronous and the second as asynchronous in our discussion.

### 3.2 Higher-order statistics confirm that astrocytes affect neuronal dynamics

To confirm the influence of glial cells on neuronal dynamics, we extend our analysis to higher-order statistics across the four case scenarios. Specifically, we examine individual cell communication using spike-train coherence (see Eq. 6). For every parameter we study, we use 1000 selected pairs from the network and verify the coherence distributions for varying numbers of glial population as shown in Fig. 8. Varying the number of glial cells elucidates the role of these elements and their communication in the frequency domain Bruns (2004); Lepage et al (2011).

**Fig 8.**
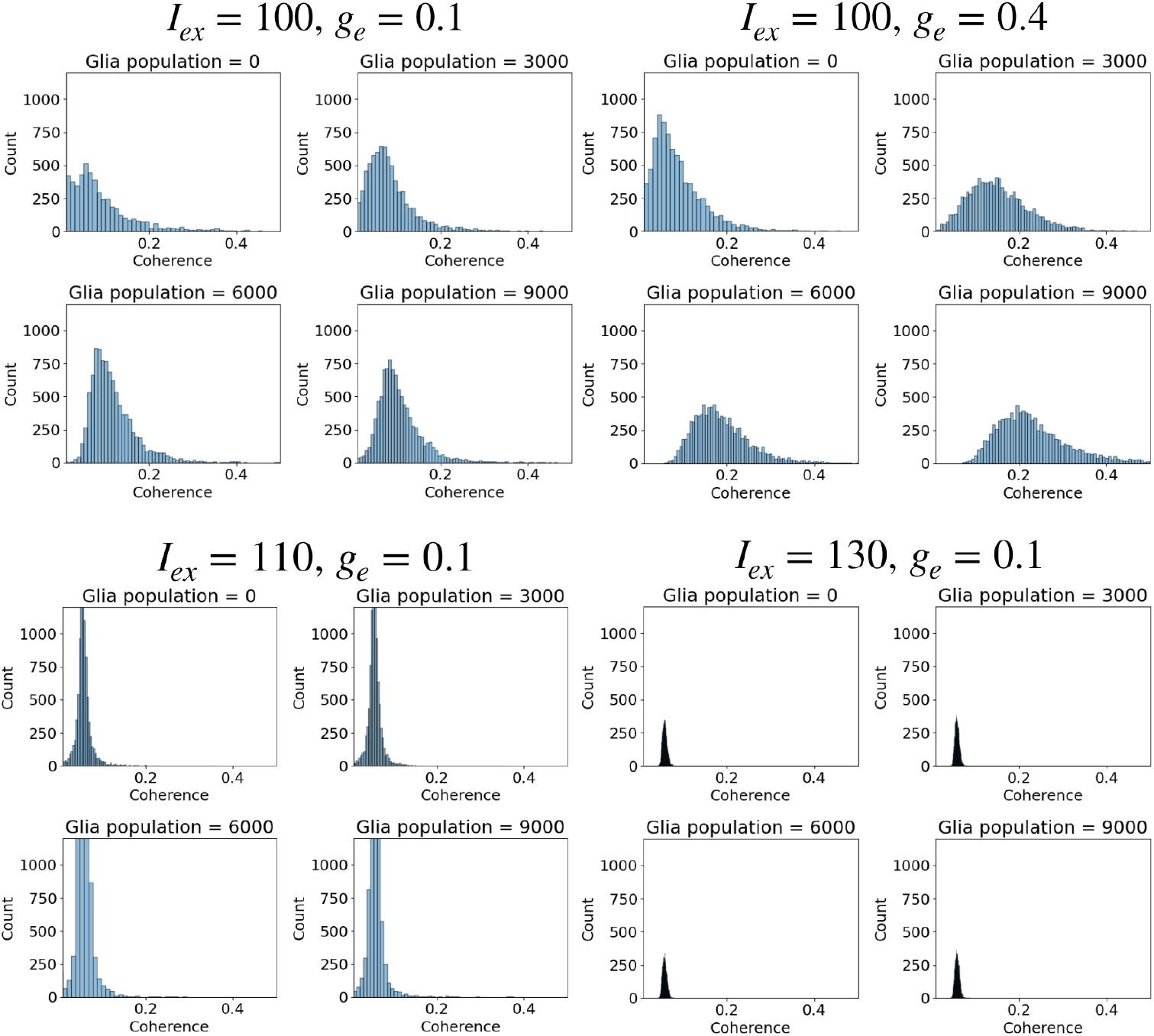
Coherence values from 1000 randomly selected neuron pairs across networks with varying glial population sizes. Baseline values are provided at the top, with parameter variations indicated in each panel title. For reference, the standard glial cell count in the network is 2000.

In Fig. 8, we observe that increasing the glial population causes a rightward, positive shift in coherence, indicating that astrocytes facilitate cells firing together at specific frequencies. This effect is particularly evident in the synchronous cases (*I*_ext_ = 100 pA, *g*_exc_ = 0.1) and (*I*_ext_ = 100 pA, *g*_exc_ = 0.4). However, the asynchronous cases (*I*_ext_ = 110 pA, *g*_exc_ = 0.1) and (*I*_ext_ = 130 pA, *g*_exc_ = 0.1) show only slight changes in coherence distribution, suggesting a weaker response. These results indicate that in synchronous conditions, glial cells enhance frequency diversity within the network. Some sample traces of individual firing rates and averages are available in Figs S1 and S2.

This set of results confirms that using the four case scenarios where glia is either present or absent with distinct activity populations is appropriate for our study of identification. In the next section, we will discuss how we use these scenarios to create a synthetic dataset for training and testing our machine learning algorithms.

### 3.3 Generation of the synthetic dataset

In order to generate a synthetic dataset in which we would train and test the different classifiers, we established four scenarios representative of distinct activity patterns: two synchronous and two asynchronous. In the synchronous cases, both used an external stimulus of *I*_ext_(*t*) = 100 pA, but with different synaptic conductance values *g*_syn_(*t*) (0.1 and 0.4). In the asynchronous cases, both had a synaptic conductance of *g*_syn_(*t*) = 0.1, but with different external stimuli *I*_ext_(*t*) (110 pA and 130 pA).

We simulated the mean firing rate across 250-time points for 1000 different network realizations. The machine learning models were then applied in three stages: first using only 50-time points, then 150, and 250-time points, to evaluate whether increasing the amount of data improves performance.

Similarly, we simulated the voltage count using 250 different voltage values across 1000 network realizations, applying the models using 50, 150, and all 250 values. This was done to investigate whether observing more voltage values improves the results. These two approaches allowed us to examine whether different recorded data collection methods (mean firing rate vs. voltage values; population and spike-based or voltage-based) produce different outcomes and whether varying levels of network synchronicity affect the quality of predictions.

To evaluate the models’ performance, the dataset was divided into 80% for training and 20% for testing following standard split in the area.

### 3.4 Data Preprocessing

For both the synchronous and asynchronous cases, using voltage count and mean firing rate as input features, we applied the Min-Max Scaler to normalize the data. This choice was primarily made to accommodate the neural networks, as the output layer of our networks used the sigmoid activation function. The sigmoid function outputs values between 0 and 1, so scaling the input features to the same range ensures better convergence and stability during the training process. This is consistent with recommendations for using the sigmoid function, as input data scaled to the activation range improves learning efficiency and model performance (LeCun et al, 2012).

However, we did not apply this scaling for the decision tree-based methods (e.g., Decision Trees, Random Forest, and Gradient Boosting), as these models are invariant to feature scaling. Decision trees split the feature space based on threshold values and do not rely on the distance between data points, which makes them unaffected by the scale of the input features. Therefore, scaling would not improve their performance, and we chose to leave the original feature scales for these models (Hastie et al, 2009).

### 3.5 Identification

In evaluating the performance of five machine learning algorithms (see Methods), we assessed their ability to identify glial cells within a test dataset. The evaluation was based on six performance metrics: accuracy, F1-Score, sensitivity, specificity, positive predictive value, and negative predictive value. Our analysis included four scenarios as we identified above. Additionally, we compared two methods of data collection: one involving mean firing rate and the other capturing voltage counts. This systematic assessment allowed us to determine which combination of algorithm and data collection method best supports the identification of glial cell influence across varying network states and stimulation intensities. We present the results for our testing data in Table 1, but see Table S1 for training results.

**Table 1.**
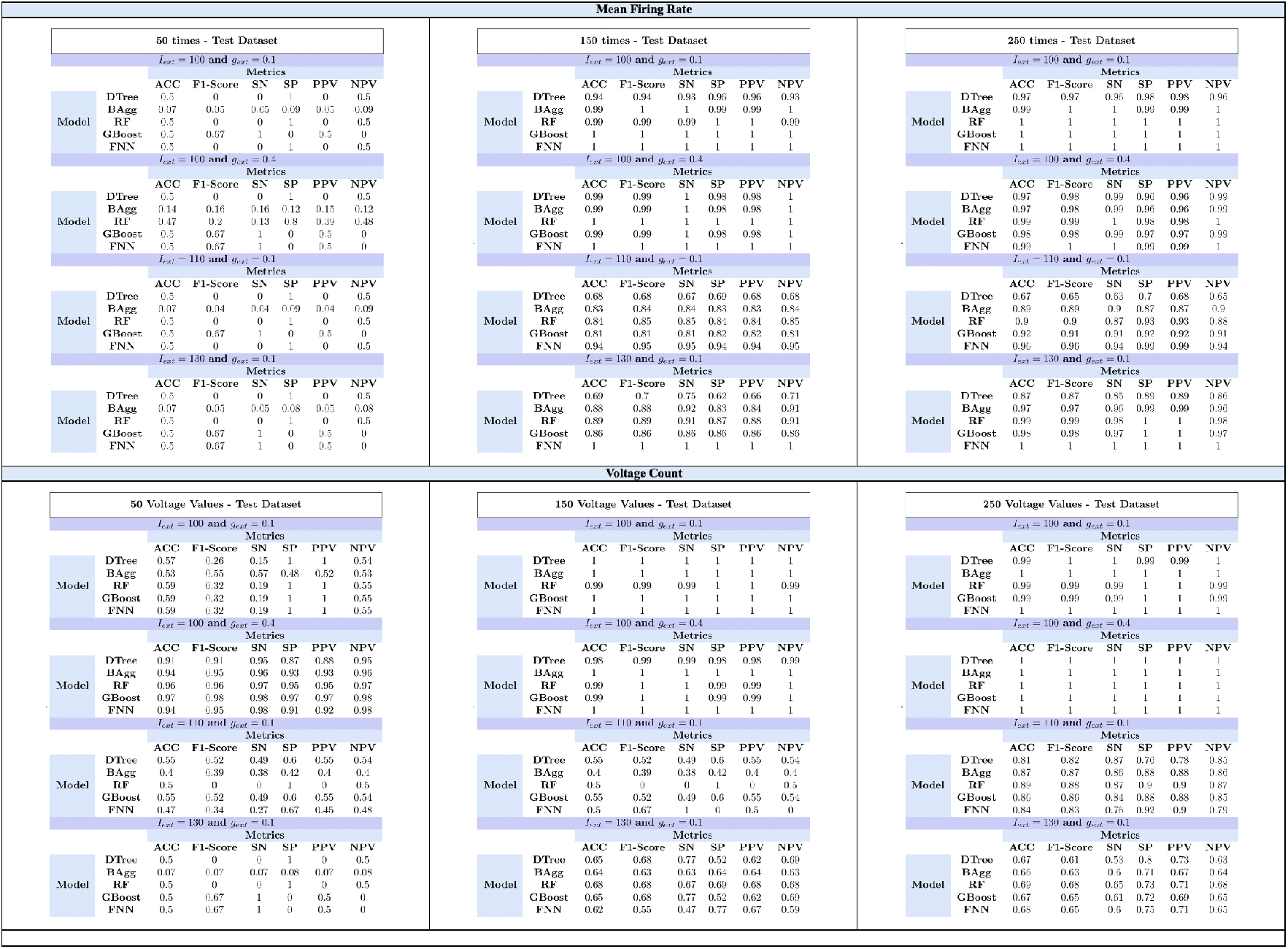
Performance of 5 machine learning algorithms (decision trees, Bagging, Random Forests, Boosting, and neural networks) in identifying glial cells in the test dataset, evaluated according to six performance measures (accuracy, F1-Score, sensitivity, specificity, positive predictive value, and negative predictive value). Two cases were considered: synchronous cases with external stimulation of 110 pA and asynchronous cases with external stimulation of 130 pA. We also investigated two different experimental collection methods: one that collects mean firing rate and the other where voltage count is obtained.

The analysis reveals that the Mean Firing Rate generally yielded superior model performance, suggesting that this brain data collection method may be more effective for identifying the presence of glial cells in neural networks. In contrast, Voltage Count achieved its best results when using 50 values, particularly in synchronous scenarios where *g*_*e*_ = 0.4. This indicates that Voltage Count-based approaches may be more suitable in synchronous cases with fewer values.

Additionally, increasing the number of predictor variables, whether time points or values in the Voltage Count, consistently improved model performance. When models were applied using Mean Firing Rate data with only 50 time points, performance was notably poor, with many models behaving as dummy classifiers. However, when the number of covariables was expanded, Feedforward Neural Networks (FNNs) out-performed other models under Mean Firing Rate in both asynchronous cases, which exhibited more complex dynamics. This reinforces the notion that asynchronous conditions are less influenced by glial cell activity, as shown by the markedly lower performance of Voltage Count models in asynchronous scenarios, even with 250 voltage values.

For Voltage Values, no model demonstrated significantly superior performance over others. However, in synchronous cases with sufficient covariables, models displayed improved accuracy, highlighting a stronger glial cell influence in these contexts.

The effectiveness of the methods used to enhance the generalization capability of the algorithms can be assessed by analyzing the performance in classifying the instances from the training and validation sets. The tables containing these results are available in the supplementary material in Table S1.

## 4 Discussions and Conclusions

Our study demonstrates the subtle yet significant role astrocytes play in modulating neuronal network dynamics, as revealed through computational simulations. By comparing network activity in scenarios with and without glial cells, we observe that astrocytes primarily influence synchronous network states, with minimal impact on asynchronous dynamics. This finding aligns with recent research suggesting that astrocytes may promote coherence in neuron firing, supporting frequency-specific synchronization within the network.

Despite these observed effects, our analysis also highlights the challenges in detecting glial cell influence using conventional metrics like firing rate and coefficient of variation (CV). While astrocytes affect network behavior in specific regimes, the changes in these common measures are often mild and subtle, making glial influence difficult to identify without high-resolution or specialized methods. Higher-order statistical approaches, such as spike-train coherence analysis, proved to be more sensitive to glial cell contributions, particularly in synchronous conditions, underscoring the value of advanced techniques in studying neuron-glia interactions.

Through spike-train coherence, we noted a distinct rightward shift in coherence distribution with increasing glial cell population, indicating that astrocytes enhance synchronized firing at specific frequencies. This effect was especially prominent in synchronous cases, where astrocytes promoted frequency diversity, suggesting a potential role in fine-tuning network oscillations. These findings open up the possibility that astrocytes may selectively influence oscillatory dynamics, contributing to network stability and information processing.

Even with the difficulties of identifying the presence of glial cells, we highlight the utility of machine learning in detecting their influence within neural networks, particularly by leveraging the Mean Firing Rate as an effective data collection method. Among the five machine learning algorithms evaluated, Feedforward Neural Networks (FNNs) demonstrated superior performance, especially in asynchronous cases characterized by complex dynamics. This aligns with our findings that asynchronous scenarios are generally less affected by glial cells, as FNNs performed best when more time points were included as predictor variables, emphasizing the importance of a richer dataset for reliable glial detection. Notably, while Mean Firing Rate consistently outperformed Voltage Count overall, the latter metric showed promise in synchronous cases with fewer values, specifically where excitatory conductance was high. This suggests that Voltage Count-based approaches may be best suited for scenarios with high coherence, while Mean Firing Rate provides a broader application across both synchronous and asynchronous conditions. Our results underscore the potential of machine learning to enhance glial cell detection, with careful consideration of data collection methods and predictor variables tailored to specific network states.

In conclusion, our computational study coupled with machine learning detection underscores the nuanced role of astrocytes in neuronal networks, particularly in synchronous regimes. By enhancing our ability to detect glial influence through advanced statistical methods, we open new avenues for exploring how neuron-glia interactions shape brain function. Future studies incorporating more complex neuron-glia models and experimental validations could further clarify the mechanisms behind astrocytic modulation, potentially contributing to the development of glia-targeted therapies or artificial intelligence systems inspired by these interactions.

## Appendix A

### Choice and initialization of parameters and hyperparameters

To evaluate the performance of each hyperparameter configuration, we used 5-fold cross-validation. This involves dividing the data into five folds, training the model on four folds, and validating it on the remaining fold. The process is repeated five times, and the mean accuracy across the folds is used to assess model performance. Cross-validation plays an important role in mitigating overfitting, as it ensures that the model generalizes well to unseen data (Kohavi, 1995). Given that our dataset is balanced, accuracy was chosen as the primary metric for model selection.

For hyperparameter tuning, we applied Bayesian Optimization using the Optuna framework (Akiba et al, 2019). Bayesian optimization, first introduced by (Mockus, 1978), is particularly efficient for navigating large and complex hyperparameter spaces, as it balances exploration and exploitation to identify optimal configurations. This made it a suitable choice for our study.

Mathematically, Bayesian optimization builds a surrogate model *p*_*surrogate*_(*y* | x), where x represents a hyperparameter configuration and *y* is the corresponding performance score (e.g., accuracy). In Optuna, we employed a Tree-structured Parzen Estimator (TPE) as the surrogate model (Bergstra et al, 2011), which models the likelihood ratio as follows:

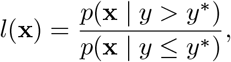

where *y*^∗^ is a threshold separating good from poor evaluations of the objective function, typically chosen based on a quantile of completed trials. By default, Optuna uses a 50% quantile, meaning trials with performance above the median are considered good.

The TPE models the two distributions *p*(x | *y > y*^∗^) and *p*(x | *y* ≤ *y*^∗^) using kernel density estimation (KDE) (Rosenblatt, 1956; Parzen, 1962) or similar techniques. In Optuna, the TPE implementation uses KDE to model the hyperparameter distributions, constructing Gaussian mixture models (Bishop, 2006a) for both good and poor trials. This allows the algorithm to estimate the likelihood ratio *l*(x) and sample new hyperparameters by maximizing this ratio, effectively focusing the search on promising regions of the hyperparameter space.

As trials progress, *y*^∗^ is dynamically updated, allowing the model to refine its search and improve efficiency. By updating the surrogate model based on each trial’s results, Bayesian optimization efficiently navigates the hyperparameter space.

We employed Optuna’s Median Pruner to reduce computational cost by halting underperforming trials early, using the following settings:

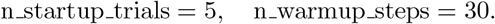

The optimization was conducted over 100 trials, with each trial representing a different configuration of hyperparameters. The combination of Bayesian optimization and cross-validation helped improve model performance while minimizing overfitting. This framework, incorporating Bayesian optimization, early pruning, and cross-validation, provided an efficient and reliable method for identifying a well-performing model for our task.

#### A.1 Decision Trees

In our optimization of the Decision Tree Classifier, we selected a range of hyperparameters that are important for controlling the complexity and performance of the model. The criterion hyperparameter, set to either *gini* or *entropy*, determines the function used to measure the quality of a split (Breiman et al, 1984b; Quinlan, 1986). The gini criterion assesses node impurity, while entropy is based on information gain.

The min samples split parameter, ranging from 2 to 50, controls the minimum number of samples required to split an internal node, helping to prevent the creation of nodes that could lead to overfitting. Setting this parameter within a broader range allows the model to consider both simpler and more complex trees. The ccp alpha parameter, which varies from 0 to 0.2, helps control tree pruning through cost complexity pruning, reducing overfitting by pruning branches that contribute little to the model (Breiman, 1984). The min samples leaf parameter, set between 1 and 50, specifies the minimum number of samples that must be present in a leaf node, ensuring that leaves contain enough samples to generalize effectively. Lastly, we defined the max depth parameter to range from 2 to 30, limiting the maximum depth of the tree and helping control the model’s complexity. This hyperparameter space was selected to allow for a balanced search for configurations that improve both model performance and generalization across different conditions.

#### A.2 Bagging

In our optimization of the Bagging Classifier, we focused on the n estimators parameter, which controls the number of base models in the ensemble. We set its range between 10 and 1000 to explore both smaller and larger ensembles. While a higher number of estimators can improve performance by reducing variance, it also increases computational cost. Our goal was to balance performance and efficiency by optimizing this parameter.

#### A.3 Random Forest

In our optimization of the Random Forest Classifier, we selected a comprehensive set of hyperparameters crucial for controlling the model’s complexity and enhancing its performance. The n estimators parameter, ranging from 100 to 1000, represents the number of trees in the forest. A larger number of estimators generally leads to improved accuracy and robustness against overfitting, as it allows the model to capture more complex patterns in the data. The max depth parameter, set between 2 and 30, limits the maximum depth of each tree, helping to prevent overfitting by restricting how deep the trees can grow. The min samples split hyperparameter, ranging from 2 to 50, determines the minimum number of samples required to split an internal node, thus avoiding the creation of nodes that may be too specific to the training data. The choice of the criterion, which can be either gini or entropy, influences the function used to measure the quality of splits, allowing the model to explore different methods of creating decision boundaries. Additionally, the ccp alpha parameter, varying from 0 to 0.1, is critical for controlling cost complexity pruning, enabling the model to prune unimportant branches and further combat overfitting. The min samples leaf parameter, set between 1 and 50, ensures that each leaf node contains a sufficient number of samples, enhancing generalization. Finally, the max features parameter allows us to specify the number of features to consider when looking for the best split, with options including *sqrt, log2*, and a fraction (0.33), promoting diversity among the trees in the forest. The hyperparameter space was designed to enable comprehensive exploration of configurations, helping to identify settings that balance performance and computational efficiency, which is important for building effective machine learning models (Breiman, 2001).

#### A.4 Gradient Boosting

In our optimization of the Gradient Boosting Classifier, we selected key hyperparameters to fine-tune model performance and manage complexity. The n estimators parameter, ranging from 100 to 1000, indicates the number of boosting stages (or trees) in the ensemble. While more estimators can reduce bias and improve accuracy, they also increase the risk of overfitting, requiring careful tuning. The max depth parameter, set between 2 and 30, limits the depth of each tree, helping to control overfitting by constraining model complexity. We included the loss hyperparameter, set to either log-loss or exponential, to specify the loss function used in optimization, allowing the model to adapt its learning strategy to the data characteristics.

The min samples split hyperparameter, ranging from 2 to 50, determines the minimum number of samples required to split an internal node, which helps regulate complexity and prevent overfitting to minor variations in the dataset. The ccp alpha parameter, varying from 0 to 0.1, controls cost complexity pruning, eliminating less informative splits to enhance generalization. The min samples leaf parameter, set between 1 and 50, ensures that each leaf node contains enough samples, further aiding generalization.

Lastly, the learning rate hyperparameter, ranging from 1 × 10^−3^ to 2 × 10^−1^, controls the contribution of each tree to the final model, adjusting how the model updates with each new tree. This hyperparameter space was designed to explore configurations that improve the Gradient Boosting Classifier’s performance while balancing accuracy and computational efficiency.

#### A.5 Feedforward Neural Networks

In our optimization of the Feedforward Neural Networks (FNNs), we focused on a variety of hyperparameters important for improving the model’s performance and ensuring effective training. The optimizer parameter plays a key role as it determines the algorithm used to update the weights during training. We included options such as *adam, rmsprop*, and *sgd* (Robbins and Monro, 1951; Hinton, 2012; Kingma and Ba, 2014) to allow the model to explore different optimization strategies, each with its strengths in handling various datasets. The activation function was chosen from *relu* (Nair and Hinton, 2010) and *tanh* (LeCun et al, 1998), with the decision made to set a single activation function for all hidden layers for simplicity, enabling a more straightforward architecture while still introducing non-linearity. We set the num layers parameter to range from 1 to 5, allowing for flexibility in the model’s depth. This range was chosen because increasing the number of layers too much can become computationally costly, while still allowing the exploration of deeper networks to capture more complex patterns in the data. For binary classification tasks, we use the sigmoid function in the output layer, which converts the linear combination into a probability value between 0 and 1 (Bishop, 2006b).

The dropout rate, ranging from 0.0 to 0.5, helps prevent overfitting by randomly dropping a fraction of neurons during training, encouraging the model to learn more robust features. The learning rate parameter, set between 1 × 10^−4^ and 1 × 10^−1^, controls the step size during weight updates, which is crucial for converging to the optimal solution effectively.

For weight initialization, we employed the *He uniform* initializer for the hidden layers using the ReLU activation function, which is particularly suited for maintaining a balanced variance of activations across layers, mitigating the vanishing gradient problem (He et al, 2015). Conversely, we utilized the *Glorot uniform* initializer for layers with the *tanh* activation function. This initializer is effective in preserving the variance of both positive and negative activations, thus enhancing the network’s ability to learn effectively without encountering issues associated with saturation in the activation function (Glorot and Bengio, 2010).

The defined hyperparameter space allows for extensive exploration and fine-tuning of the FNN architecture, optimizing generalization to unseen data while balancing computational efficiency.

## Acknowledgments

J.P.P. was supported by grants #2023*/*09094 − 0 and #2023*/*15585 − 6, São Paulo Research Foundation (FAPESP). P.R.P. was supported by grant #2020*/*04624 − 2, São Paulo Research Foundation (FAPESP). R.F.O.P. was supported by the Palm Health-Sponsored Program in Computational Brain Science and Health, FAU Stiles-Nicholson Brain Institute. R.F.O.P and L.F. are supported by the Jupiter Life Science Initiative (JLSI).

## References

Adamczyk A (2023) Glial–neuronal interactions in neurological disorders: Molecular mechanisms and potential points for intervention. International Journal of Molecular Sciences 24(7):6274

Akiba T, Sano S, Yanase T, et al (2019) Optuna: A next-generation hyperparameter optimization framework. In: Proceedings of the 25th ACM SIGKDD International Conference on Knowledge Discovery & Data Mining, pp 2623–2631

Bayraktar OA, Fuentealba LC, Alvarez-Buylla A, et al (2015) Astrocyte development and heterogeneity. Cold Spring Harbor perspectives in biology 7(1):a020362

Bazargani N, Attwell D (2016) Astrocyte calcium signaling: the third wave. Nature Neuroscience 19(2):182–189

Ben Achour S, Pascual O (2012) Astrocyte–neuron communication: functional consequences. Neurochemical research 37:2464–2473

Bergstra J, Bardenet R, Bengio Y, et al (2011) Algorithms for hyper-parameter optimization. In: Advances in Neural Information Processing Systems, pp 2546–2554

Birch AM (2014) The contribution of astrocytes to alzheimer’s disease. Biochemical Society Transactions 42(5):1316–1320

Bishop CM (2006a) Pattern recognition and machine learning. Springer Science & Business Media

Bishop CM (2006b) Pattern recognition and machine learning. Springer

Borges F, Protachevicz PR, Pena R, et al (2020) Self-sustained activity of low firing rate in balanced networks. Physica A: Statistical Mechanics and its Applications 537:122671

Breiman L (1984) Classification and regression trees (CART). Wadsworth International Group, Belmont, California

Breiman L (1996) Bagging predictors. Machine Learning 24(2):123–140

Breiman L (2001) Random forests. Machine learning 45(1):5–32

Breiman L, Friedman JH, Olshen RA, et al (1984a) Classification and Regression Trees. Wadsworth International Group, Belmont, CA

Breiman L, Friedman JH, Stone CJ, et al (1984b) Classification and Regression Trees. CRC Press

Bruns A (2004) Fourier-, hilbert-and wavelet-based signal analysis: are they really different approaches? Journal of neuroscience methods 137(2):321–332

Burkitt A (2006) A review of the integrate-and-fire neuron model: I. homogeneous synaptic input. Biological cybernetics 95:1–19

Cai Z, Wan CQ, Liu Z (2017) Astrocyte and alzheimer’s disease. Journal of neurology 264:2068–2074

Cessac B, Viéville T (2008) On dynamics of integrate-and-fire neural networks with conductance based synapses. Frontiers in computational neuroscience 2:228

Chen J, Poskanzer KE, Freeman MR, et al (2020) Live-imaging of astrocyte morphogenesis and function in zebrafish neural circuits. Nature neuroscience 23(10):1297–1306

Covelo A, Araque A (2018) Neuronal activity determines distinct gliotransmitter release from a single astrocyte. eLife 7:e32237

De Pittá M, Brunel N (2022) Multiple forms of working memory emerge from synapse– astrocyte interactions in a neuron–glia network model. Proceedings of the National Academy of Sciences 119(43):e2207912119

Destexhe A (1997) Conductance-based integrate-and-fire models. Neural Computation 9(3):503–514

Dimou L, Götz M (2014) Glial cells as progenitors and stem cells: New roles in the healthy and diseased brain. Physiological Reviews 94(3):709–737

Fellin T (2009) Communication between neurons and astrocytes: relevance to the modulation of synaptic and network activity. Journal of Neurochemistry 108(3):533– 544

Friedman JH (2001) Greedy function approximation: a gradient boosting machine. Annals of Statistics pp 1189–1232

Gerstner W, Kistler WM, Naud R, et al (2014) Neuronal dynamics: From single neurons to networks and models of cognition. Cambridge University Press

Gini C (1912) Variabilitá e mutabilitá: contributo allo studio delle distribuzioni e delle relazioni statistiche.[Fasc. I.]. Tipogr. di P. Cuppini

Glorot X, Bengio Y (2010) Understanding the difficulty of training deep feedforward neural networks. AISTATS 9:249–256

Gómez-Gonzalo M, Navarrete M, Perea G, et al (2015) Endocannabinoids induce lateral long-term potentiation of transmitter release by stimulation of gliotransmission. Cerebral cortex 25(10):3699–3712

Guerra-Gomes S, Sousa N, Pinto L, et al (2018) Functional roles of astrocyte calcium elevations: from synapses to behavior. Frontiers in cellular neuroscience 11:427

Halassa MM, Haydon PG (2010) Integrated brain circuits: astrocytic networks modulate neuronal activity and behavior. Annual Review of Physiology 72(1):335–355

Handy G, Borisyuk A (2023) Investigating the ability of astrocytes to drive neural network synchrony. PLoS computational biology 19(8):e1011290

Hastie T, Tibshirani R, Friedman J (2009) The Elements of Statistical Learning: Data Mining, Inference, and Prediction. Springer

He F, Sun YE (2007) Glial cells more than support cells? The international journal of biochemistry & cell biology 39(4):661–665

He K, Zhang X, Ren S, et al (2015) Delving deep into rectifiers: Surpassing human-level performance on imagenet classification. arXiv preprint 150201852

Hinton G (2012) Lecture 6e rmsprop: Divide the gradient by a running average of its recent magnitude. Coursera: Neural Networks for Machine Learning

Ian Goodfellow ACYoshua Bengio (2016) Deep learning. MIT press

Jackson FR (2011) Glial cell modulation of circadian rhythms. Glia 59(9):1341–1350

Jäkel S, Dimou L (2017) Glial cells and their function in the adult brain: a journey through the history of their ablation. Frontiers in cellular neuroscience 11:24

Jessen KR (2004) Glial cells. The international journal of biochemistry & cell biology 36(10):1861–1867

Jäkel S, Dimou L (2017) Glial cells and their function in the adult brain: A journey through the history of their ablation. Frontiers in Cellular Neuroscience 11

Kanner S, Goldin M, Galron R, et al (2018) Astrocytes restore connectivity and synchronization in dysfunctional cerebellar networks. Proceedings of the National Academy of Sciences 115(31):8025–8030

Kim NS, Chung WS (2023) Astrocytes regulate neuronal network activity by mediating synapse remodeling. Neuroscience Research 187:3–13

Kingma DP, Ba J (2014) Adam: A method for stochastic optimization. arXiv preprint 14126980

Kohavi R (1995) A study of cross-validation and bootstrap for accuracy estimation and model selection. In: Proceedings of the 14th International Joint Conference on Artificial Intelligence (IJCAI), pp 1137–1145

Kol A, Adamsky A, Groysman M, et al (2020) Astrocytes contribute to remote memory formation by modulating hippocampal–cortical communication during learning. Nature neuroscience 23(10):1229–1239

Lawal O, Ulloa Severino FP, Eroglu C (2022) The role of astrocyte structural plasticity in regulating neural circuit function and behavior. Glia 70(8):1467–1483

LeCun Y, Bottou L, Bengio Y, et al (1998) Gradient-based learning applied to document recognition. Proceedings of the IEEE 86(11):2278–2324

LeCun Y, Bottou L, Orr GB, et al (2012) Efficient BackProp. Springer

Lengler J, Steger A (2017) Note on the coefficient of variations of neuronal spike trains. Biological Cybernetics 111

Lepage KQ, Kramer MA, Eden UT (2011) The dependence of spike field coherence on expected intensity. Neural computation 23(9):2209–2241

Li YX, Rinzel J (1994) Equations for insp3 receptor-mediated [ca2+] i oscillations derived from a detailed kinetic model: a hodgkin-huxley like formalism. Journal of theoretical Biology 166(4):461–473

Lines J, Martin ED, Kofuji P, et al (2020) Astrocytes modulate sensory-evoked neuronal network activity. Nature communications 11(1):3689

Linne ML, Aćimović J, Saudargiene A, et al (2022) Neuron–glia interactions and brain circuits pp 87–103

Liu X, Ying J, Wang X, et al (2021) Astrocytes in neural circuits: key factors in synaptic regulation and potential targets for neurodevelopmental disorders. Frontiers in Molecular Neuroscience 14:729273

McCulloch WS, Pitts W (1943) A logical calculus of the ideas immanent in nervous activity. The Bulletin of Mathematical Biophysics 5:115–133

Mockus J (1978) Application of bayesian approach to numerical methods of global and stochastic optimization. Journal of Global Optimization 4:173–190

Nair V, Hinton GE (2010) Rectified linear units improve restricted boltzmann machines. In: Proceedings of the 27th International Conference on Machine Learning (ICML-10), pp 807–814

Newman EA (2015) Glial cell regulation of neuronal activity and blood flow in the retina by release of gliotransmitters. Philosophical Transactions of the Royal Society B: Biological Sciences 370(1672):20140195

Panatier A, Vallée J, Haber M, et al (2011) Astrocytes are endogenous regulators of basal transmission at central synapses. Cell 146(5):785–798

Paolicelli RC, Bolasco G, Pagani F, et al (2011) Synaptic pruning by microglia is necessary for normal brain development. science 333(6048):1456–1458

Parpura V, Heneka MT, Montana V, et al (2012) Glial cells in (patho) physiology. Journal of neurochemistry 121(1):4–27

Parzen E (1962) On estimation of a probability density function and mode. The Annals of Mathematical Statistics 33(3):1065–1076

Pena RF, Vellmer S, Bernardi D, et al (2018) Self-consistent scheme for spiketrain power spectra in heterogeneous sparse networks. Frontiers in computational neuroscience 12:9

Perea G, Araque A (2007) Astrocytes potentiate transmitter release at single hippocampal synapses. Science 317(5841):1083–1086

Perez-Catalan NA, Doe CQ, Ackerman SD (2021) The role of astrocyte-mediated plasticity in neural circuit development and function. Neural development 16(1):1

Poskanzer KE, Yuste R (2016) Astrocytes regulate cortical state switching in vivo. Proceedings of the National Academy of Sciences 113(19):E2675–E2684

Protachevicz PR, Borges FS, Iarosz KC, et al (2020) Influence of delayed conductance on neuronal synchronization. Frontiers in Physiology 11:1053

Protachevicz PR, Bonin C, Iarosz K, et al (2022) Large coefficient of variation of inter-spike intervals induced by noise current in the resonate-and-fire model neuron. Cognitive Neurodynamics 16(6):1461–1470

Purushotham SS, Buskila Y (2023) Astrocytic modulation of neuronal signalling. Frontiers in Network Physiology 3:1205544

Quinlan JR (1986) Induction of decision trees. Machine Learning 1(1):81–106

Robbins H, Monro S (1951) A stochastic approximation method. The Annals of Mathematical Statistics 22(3):400–407

Rosenblatt F (1958) The perceptron: a probabilistic model for information storage and organization in the brain. Psychological Review 65(6):386

Rosenblatt M (1956) Remarks on some nonparametric estimates of a density function. The Annals of Mathematical Statistics 27(3):832–837

Sanaullah, Koravuna S, Rückert U, et al (2023) Exploring spiking neural networks: a comprehensive analysis of mathematical models and applications. Frontiers in Computational Neuroscience 17:1215824

Sanz-Gálvez R, Falardeau D, Kolta A, et al (2024) The role of astrocytes from synaptic to non-synaptic plasticity. Frontiers in Cellular Neuroscience 18:1477985

Shannon CE (1948) A mathematical theory of communication. The Bell System Technical Journal 27(3):379–423. 10.1002/j.1538-7305.1948.tb01338.x

Sherwood MW, Arizono M, Hisatsune C, et al (2017) Astrocytic ip3rs: Contribution to ca2+ signalling and hippocampal ltp. Glia 65(3):502–513

Sherwood MW, Arizono M, Panatier A, et al (2021) Astrocytic ip3rs: Beyond ip3r2. Frontiers in cellular neuroscience 15:695817

Shimoura RO, Pena RF, Lima V, et al (2021) Building a model of the brain: from detailed connectivity maps to network organization. The European Physical Journal Special Topics 230(14):2887–2909

Stackhouse TL, Mishra A (2021) Neurovascular coupling in development and disease: focus on astrocytes. Frontiers in cell and developmental biology 9:702832

Stevens B, Allen NJ, Vazquez LE, et al (2007) The classical complement cascade mediates cns synapse elimination. Cell 131(6):1164–1178

Stimberg M, Goodman DF, Brette R, et al (2019) Modeling neuron–glia interactions with the brian 2 simulator. Computational glioscience pp 471–505

Straub SV, Nelson MT (2007) Astrocytic calcium signaling: the information currency coupling neuronal activity to the cerebral microcirculation. Trends in cardiovascular medicine 17(6):183–190

Tomov P, Pena RFO, Roque AC, et al (2016) Mechanisms of self-sustained oscillatory states in hierarchical modular networks with mixtures of electrophysiological cell types. Frontiers in Computational Neuroscience 10

Van Horn MR, Benfey NJ, Shikany C, et al (2021) Neuron-astrocyte networking: astrocytes orchestrate and respond to changes in neuronal network activity across brain states and behaviors. Journal of neurophysiology 126(2):627–636

Von Bartheld CS, Bahney J, Herculano-Houzel S (2016) The search for true numbers of neurons and glial cells in the human brain: A review of 150 years of cell counting. Journal of Comparative Neurology 524(18):3865–3895

Wang F, Xu Q, Wang W, et al (2012) Bergmann glia modulate cerebellar purkinje cell bistability via ca2+-dependent k+ uptake. Proceedings of the National Academy of Sciences 109(20):7911–7916

Wei F, Shuai J (2011) Intercellular calcium waves in glial cells with bistable dynamics. Physical biology 8(2):026009

Wu J, Chen XY, Zhang H, et al (2019) Hyperparameter optimization for machine learning models based on bayesian optimization. Journal of Electronic Science and Technology 17(1):26–40

