## Supporting Material for "Astrocytic Signatures in Neuronal Activity: A Machine Learning-Based Identification Approach"

**Table S1**: Performance of 5 machine learning algorithms (decision trees, Bagging, Random Forests, Boosting, and neural networks) in identifying glial cells in the train dataset, evaluated according to six performance measures (accuracy, F1-Score, sensitivity, specificity, positive predictive value, and negative predictive value). Two cases were considered: synchronous cases with external stimulation of 110 pA and asynchronous cases with external stimulation of 130 pA. We also investigated two different experimental collection methods: one that collects mean firing rate and the other where voltage count is obtained.

**Mean Firing Rate**

**50 times - Train Dataset**

| Condition | Model | ACC | F1-Score | SN | SP | PPV | NPV |
|---|---|---|---|---|---|---|---|
| $I_{ext} = 100$ and $g_{ext} = 0.1$ | DTree | 0.5 | 0 | 0 | 1 | 0 | 0.5 |
| | BAgg | 0.95 | 0.95 | 0.95 | 0.95 | 0.95 | 0.95 |
| | RF | 0.5 | 0 | 0 | 1 | 0 | 0.5 |
| | GBoost | 0.5 | 0.67 | 1 | 0 | 0.5 | 0 |
| | FNN | 0.5 | 0 | 0 | 1 | 0 | 0.5 |
| $I_{ext} = 100$ and $g_{ext} = 0.4$ | DTree | 0.5 | 0 | 0 | 1 | 0 | 0.5 |
| | BAgg | 0.93 | 0.93 | 0.92 | 0.94 | 0.94 | 0.92 |
| | RF | 0.52 | 0.27 | 0.18 | 0.87 | 0.58 | 0.51 |
| | GBoost | 0.5 | 0.67 | 1 | 0 | 0.5 | 0 |
| | FNN | 0.5 | 0.67 | 1 | 0 | 0.5 | 0 |
| $I_{ext} = 110$ and $g_{ext} = 0.1$ | DTree | 0.5 | 0 | 0 | 1 | 0 | 0.5 |
| | BAgg | 0.81 | 0.81 | 0.81 | 0.81 | 0.81 | 0.81 |
| | RF | 0.5 | 0 | 0 | 1 | 0 | 0.5 |
| | GBoost | 0.5 | 0.67 | 1 | 0 | 0.5 | 0 |
| | FNN | 0.5 | 0 | 0 | 1 | 0 | 0.5 |
| $I_{ext} = 130$ and $g_{ext} = 0.1$ | DTree | 0.5 | 0 | 0 | 1 | 0 | 0.5 |
| | BAgg | 0.62 | 0.63 | 0.64 | 0.61 | 0.62 | 0.63 |
| | RF | 0.5 | 0 | 0 | 1 | 0 | 0.5 |
| | GBoost | 0.5 | 0.67 | 1 | 0 | 0.5 | 0 |
| | FNN | 0.5 | 0.67 | 1 | 0 | 0.5 | 0 |

**150 times - Train Dataset**

| Condition | Model | ACC | F1-Score | SN | SP | PPV | NPV |
|---|---|---|---|---|---|---|---|
| $I_{ext} = 100$ and $g_{ext} = 0.1$ | DTree | 0.98 | 0.99 | 0.99 | 0.98 | 0.98 | 0.99 |
| | BAgg | 1 | 1 | 1 | 1 | 1 | 1 |
| | RF | 1 | 1 | 1 | 1 | 1 | 1 |
| | GBoost | 1 | 1 | 1 | 1 | 1 | 1 |
| | FNN | 1 | 1 | 1 | 1 | 1 | 1 |
| $I_{ext} = 100$ and $g_{ext} = 0.4$ | DTree | 1 | 1 | 1 | 1 | 1 | 1 |
| | BAgg | 1 | 1 | 1 | 1 | 1 | 1 |
| | RF | 1 | 1 | 1 | 1 | 1 | 1 |
| | GBoost | 1 | 1 | 1 | 1 | 1 | 1 |
| | FNN | 1 | 1 | 1 | 1 | 1 | 1 |
| $I_{ext} = 110$ and $g_{ext} = 0.1$ | DTree | 0.82 | 0.82 | 0.79 | 0.86 | 0.85 | 0.8 |
| | BAgg | 1 | 1 | 1 | 1 | 1 | 1 |
| | RF | 0.98 | 0.98 | 0.98 | 0.98 | 0.98 | 0.98 |
| | GBoost | 1 | 1 | 1 | 1 | 1 | 1 |
| | FNN | 0.99 | 0.99 | 0.98 | 1 | 1 | 0.99 |
| $I_{ext} = 130$ and $g_{ext} = 0.1$ | DTree | 0.85 | 0.85 | 0.86 | 0.83 | 0.84 | 0.86 |
| | BAgg | 1 | 1 | 1 | 1 | 1 | 1 |
| | RF | 1 | 1 | 1 | 1 | 1 | 1 |
| | GBoost | 1 | 1 | 1 | 1 | 1 | 1 |
| | FNN | 1 | 1 | 1 | 1 | 1 | 1 |

**250 times - Train Dataset**

| Condition | Model | ACC | F1-Score | SN | SP | PPV | NPV |
|---|---|---|---|---|---|---|---|
| $I_{ext} = 100$ and $g_{ext} = 0.1$ | DTree | 0.99 | 0.99 | 0.99 | 0.98 | 0.98 | 0.99 |
| | BAgg | 1 | 1 | 1 | 1 | 1 | 1 |
| | RF | 1 | 1 | 1 | 1 | 1 | 1 |
| | GBoost | 1 | 1 | 1 | 1 | 1 | 1 |
| | FNN | 1 | 1 | 1 | 1 | 1 | 1 |
| $I_{ext} = 100$ and $g_{ext} = 0.4$ | DTree | 1 | 1 | 1 | 0.99 | 1 | 1 |
| | BAgg | 1 | 1 | 1 | 1 | 1 | 1 |
| | RF | 1 | 1 | 1 | 1 | 1 | 1 |
| | GBoost | 1 | 1 | 1 | 1 | 1 | 1 |
| | FNN | 1 | 1 | 1 | 1 | 1 | 1 |
| $I_{ext} = 110$ and $g_{ext} = 0.1$ | DTree | 0.88 | 0.88 | 0.86 | 0.9 | 0.89 | 0.87 |
| | BAgg | 1 | 1 | 1 | 1 | 1 | 1 |
| | RF | 1 | 1 | 1 | 1 | 1 | 1 |
| | GBoost | 1 | 1 | 1 | 1 | 1 | 1 |
| | FNN | 1 | 1 | 1 | 1 | 1 | 1 |
| $I_{ext} = 130$ and $g_{ext} = 0.1$ | DTree | 0.95 | 0.95 | 0.93 | 0.97 | 0.97 | 0.93 |
| | BAgg | 1 | 1 | 1 | 1 | 1 | 1 |
| | RF | 1 | 1 | 1 | 1 | 1 | 1 |
| | GBoost | 1 | 1 | 1 | 1 | 1 | 1 |
| | FNN | 1 | 1 | 1 | 1 | 1 | 1 |

**Voltage Count**

**50 Voltage Values - Train Dataset**

| Condition | Model | ACC | F1-Score | SN | SP | PPV | NPV |
|---|---|---|---|---|---|---|---|
| $I_{ext} = 100$ and $g_{ext} = 0.1$ | DTree | 0.56 | 0.23 | 0.13 | 1 | 1 | 0.53 |
| | BAgg | 0.72 | 0.71 | 0.68 | 0.76 | 0.74 | 0.7 |
| | RF | 0.57 | 0.25 | 0.14 | 1 | 1 | 0.54 |
| | GBoost | 0.57 | 0.25 | 0.14 | 1 | 1 | 0.54 |
| | FNN | 0.57 | 0.25 | 0.14 | 1 | 1 | 0.54 |
| $I_{ext} = 100$ and $g_{ext} = 0.4$ | DTree | 0.98 | 0.98 | 0.97 | 0.99 | 0.99 | 0.97 |
| | BAgg | 1 | 1 | 1 | 1 | 1 | 1 |
| | RF | 1 | 1 | 1 | 1 | 1 | 1 |
| | GBoost | 1 | 1 | 1 | 1 | 1 | 1 |
| | FNN | 0.99 | 0.99 | 0.99 | 0.99 | 0.99 | 0.99 |
| $I_{ext} = 110$ and $g_{ext} = 0.1$ | DTree | 0.56 | 0.51 | 0.47 | 0.65 | 0.57 | 0.55 |
| | BAgg | 0.93 | 0.92 | 0.92 | 0.94 | 0.93 | 0.92 |
| | RF | 0.5 | 0 | 0 | 1 | 0 | 0.5 |
| | GBoost | 0.56 | 0.51 | 0.47 | 0.65 | 0.57 | 0.55 |
| | FNN | 0.59 | 0.48 | 0.38 | 0.8 | 0.66 | 0.56 |
| $I_{ext} = 130$ and $g_{ext} = 0.1$ | DTree | 0.5 | 0 | 0 | 1 | 0 | 0.5 |
| | BAgg | 0.61 | 0.62 | 0.64 | 0.57 | 0.6 | 0.61 |
| | RF | 0.5 | 0 | 0 | 1 | 0 | 0.5 |
| | GBoost | 0.5 | 0.67 | 1 | 0 | 0.5 | 0 |
| | FNN | 0.5 | 0.67 | 1 | 0 | 0.5 | 0 |

**150 Voltage Values - Train Dataset**

| Condition | Model | ACC | F1-Score | SN | SP | PPV | NPV |
|---|---|---|---|---|---|---|---|
| $I_{ext} = 100$ and $g_{ext} = 0.1$ | DTree | 1 | 1 | 1 | 0.99 | 0.99 | 1 |
| | BAgg | 1 | 1 | 1 | 1 | 1 | 1 |
| | RF | 1 | 1 | 0.99 | 1 | 1 | 1 |
| | GBoost | 1 | 1 | 1 | 1 | 1 | 1 |
| | FNN | 1 | 1 | 1 | 1 | 1 | 1 |
| $I_{ext} = 100$ and $g_{ext} = 0.4$ | DTree | 1 | 1 | 1 | 0.99 | 0.99 | 1 |
| | BAgg | 1 | 1 | 1 | 1 | 1 | 1 |
| | RF | 1 | 1 | 1 | 0.99 | 1 | 1 |
| | GBoost | 1 | 1 | 1 | 1 | 1 | 1 |
| | FNN | 1 | 1 | 1 | 1 | 1 | 1 |
| $I_{ext} = 110$ and $g_{ext} = 0.1$ | DTree | 0.56 | 0.51 | 0.47 | 0.65 | 0.57 | 0.55 |
| | BAgg | 0.96 | 0.96 | 0.95 | 0.96 | 0.96 | 0.96 |
| | RF | 0.5 | 0 | 0 | 1 | 0 | 0.5 |
| | GBoost | 0.56 | 0.51 | 0.47 | 0.65 | 0.57 | 0.55 |
| | FNN | 0.5 | 0.67 | 1 | 0 | 0.5 | 0 |
| $I_{ext} = 130$ and $g_{ext} = 0.1$ | DTree | 0.64 | 0.7 | 0.82 | 0.47 | 0.61 | 0.72 |
| | BAgg | 1 | 1 | 1 | 1 | 1 | 1 |
| | RF | 0.69 | 0.7 | 0.72 | 0.66 | 0.68 | 0.7 |
| | GBoost | 0.64 | 0.7 | 0.82 | 0.47 | 0.61 | 0.72 |
| | FNN | 0.62 | 0.59 | 0.55 | 0.69 | 0.64 | 0.61 |

**250 Voltage Values - Train Dataset**

| Condition | Model | ACC | F1-Score | SN | SP | PPV | NPV |
|---|---|---|---|---|---|---|---|
| $I_{ext} = 100$ and $g_{ext} = 0.1$ | DTree | 1 | 1 | 1 | 0.99 | 0.99 | 1 |
| | BAgg | 1 | 1 | 1 | 1 | 1 | 1 |
| | RF | 1 | 1 | 1 | 1 | 1 | 1 |
| | GBoost | 1 | 1 | 1 | 1 | 1 | 1 |
| | FNN | 1 | 1 | 1 | 1 | 1 | 1 |
| $I_{ext} = 100$ and $g_{ext} = 0.4$ | DTree | 1 | 1 | 1 | 1 | 1 | 1 |
| | BAgg | 1 | 1 | 1 | 1 | 1 | 1 |
| | RF | 1 | 1 | 1 | 1 | 1 | 1 |
| | GBoost | 1 | 1 | 1 | 1 | 1 | 1 |
| | FNN | 1 | 1 | 1 | 1 | 1 | 1 |
| $I_{ext} = 110$ and $g_{ext} = 0.1$ | DTree | 0.87 | 0.88 | 0.91 | 0.83 | 0.85 | 0.91 |
| | BAgg | 1 | 1 | 1 | 1 | 1 | 1 |
| | RF | 0.95 | 0.95 | 0.94 | 0.96 | 0.96 | 0.94 |
| | GBoost | 0.96 | 0.96 | 0.96 | 0.97 | 0.97 | 0.96 |
| | FNN | 0.94 | 0.94 | 0.88 | 1 | 1 | 0.89 |
| $I_{ext} = 130$ and $g_{ext} = 0.1$ | DTree | 0.7 | 0.67 | 0.62 | 0.78 | 0.74 | 0.67 |
| | BAgg | 1 | 1 | 1 | 1 | 1 | 1 |
| | RF | 0.84 | 0.84 | 0.84 | 0.84 | 0.84 | 0.84 |
| | GBoost | 0.72 | 0.72 | 0.72 | 0.71 | 0.72 | 0.72 |
| | FNN | 0.73 | 0.72 | 0.69 | 0.77 | 0.75 | 0.71 |

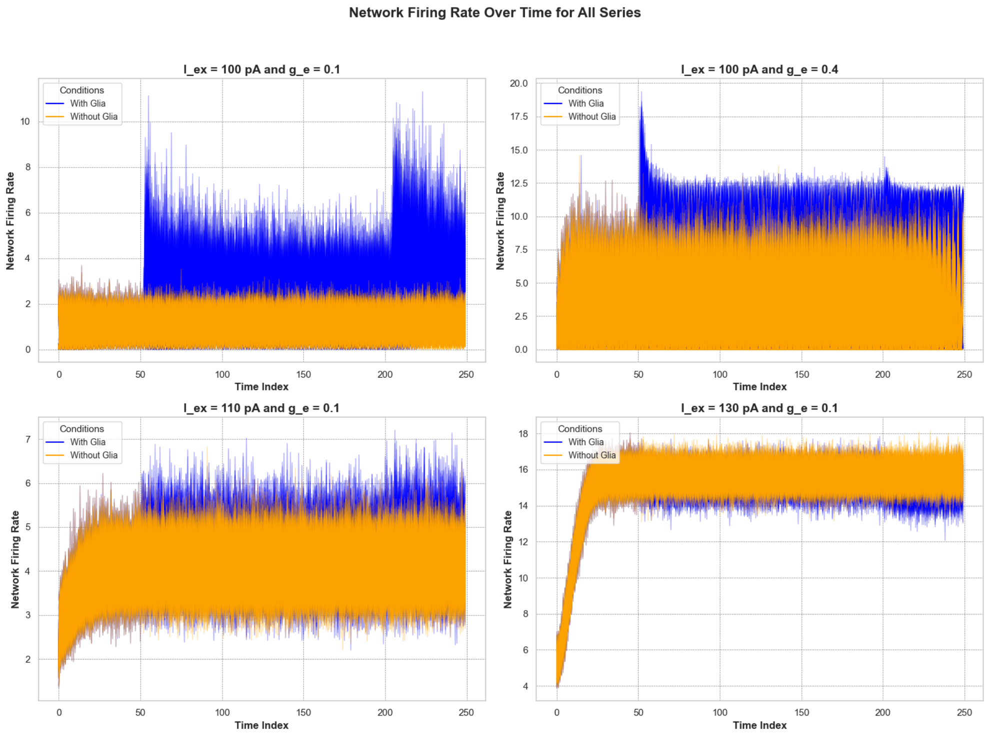


**Fig. S1**: Sample traces for network firing rate for all the time series considering all 4 scenarios that were proposed: ($I_{\text{ext}} = 100$ pA, $g_{ex} = 0.1$), ($I_{ex} = 100$ pA, $g_{ex} = 0.4$), ($I_{ex} = 110$ pA, $g_{ex} = 0.1$), and ($I_{ex} = 130$ pA, $g_{ex} = 0.1$). Blue: with glial cells. Yellow: without glial cells.

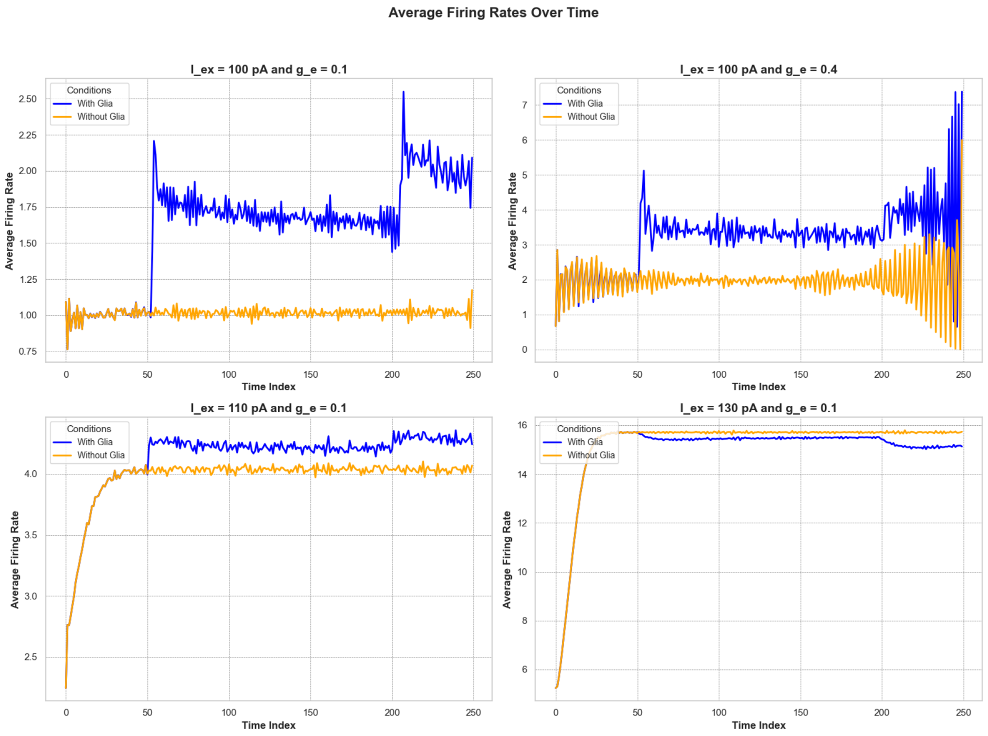


**Fig. S2**: Sample traces for network averaged firing rate over all the time series considering all 4 scenarios that were proposed: ($I_{\text{ext}} = 100$ pA, $g_{ex} = 0.1$), ($I_{ex} = 100$ pA, $g_{ex} = 0.4$), ($I_{ex} = 110$ pA, $g_{ex} = 0.1$), and ($I_{ex} = 130$ pA, $g_{ex} = 0.1$). Blue: with glial cells. Yellow: without glial cells.

| Parameter | Value | Description |
|---|---|---|
| stimulus | 1.2 | Stimulus strength |
| $g_e$ | 0.05 | Input excitatory synaptic conductance scaling |
| $g_i$ | 1.0 | Input inhibitory synaptic conductance scaling |
| $\rho_{e_{input}}$ | 6.5e-4 | Astrocytic vesicle-to-extracellular volume ratio |
| UA_input | 0.6 | Gliotransmitter release probability |
| G_T_input | 200 | Total vesicular gliotransmitter concentration |
| C_thetainput | 0.5 | $Ca^{2+}$ threshold for exocytosis |
| Nglia | 2000 | Number of astrocytes |
| N_e | 1000 | Number of excitatory neurons |
| N_i | 250 | Number of inhibitory neurons |
| N_a | Nglia | Number of astrocytes |
| size | 3.75 mm | Size of square lattice |
| distance | 50 µm | Distance between neurons |
| E_l | -60 mV | Leak reversal potential |
| g_l | 9.99 nS | Leak conductance |
| E_e | 0 mV | Excitatory synaptic reversal potential |
| E_i | -80 mV | Inhibitory synaptic reversal potential |
| C_m | 198 pF | Membrane capacitance |
| tau_e | 5 ms | Excitatory synaptic time constant |
| tau_i | 10 ms | Inhibitory synaptic time constant |
| tau_r | 5 ms | Refractory period |
| I_app | 100 pA | External current |
| V_th | -50 mV | Firing threshold |
| V_r | E_l | Reset potential |
| rho_c | 0.005 | Synaptic vesicle-to-extracellular space volume ratio |
| Y_T | 500 mM | Total vesicular neurotransmitter concentration |
| Omega_c | 40 /s | Neurotransmitter clearance rate |
| U_0__star | 0.6 | Resting synaptic release probability |
| Omega_f | 3.33 /s | Synaptic facilitation rate |
| Omega_d | 2.0 /s | Synaptic depression rate |
| O_G | 1.5 /µM/s | Agonist binding rate |
| Omega_G | 0.5 /60 s | Agonist release rate |
| O_P | 0.9 µM/s | Maximal $Ca^{2+}$ uptake rate by SERCAs |
| K_P | 0.05 µM | $Ca^{2+}$ affinity of SERCAs |
| C_T | 2 µM | Total cell free $Ca^{2+}$ content |
| rho_A | 0.18 | ER-to-cytoplasm volume ratio |
| Omega_C | 6 /s | Maximal rate of $Ca^{2+}$ release by IPRs |
| Omega_L | 0.1 /s | Maximal rate of $Ca^{2+}$ leak from the ER |
| d_1 | 0.13 µM | $IP_3$ binding affinity |
| d_2 | 1.05 µM | $Ca^{2+}$ inactivation dissociation constant |
| O_2 | 0.2 /µM/s | IPR binding rate for $Ca^{2+}$ inhibition |
| d_3 | 0.9434 µM | IP dissociation constant |
| d_5 | 0.08 µM | $Ca^{2+}$ activation dissociation constant |
| O_beta | 0.5 µM/s | Maximal rate of $IP_3$ production by PLCbeta |
| O_N | 0.3 /µM/s | Agonist binding rate |
| Omega_N | 0.5 /s | Maximal inactivation rate |
| K_KC | 0.5 µM | $Ca^{2+}$ affinity of PKC |
| zeta | 10 | Maximal reduction of receptor affinity by PKC |
| O_delta | 4.8 µM/s | Maximal rate of $IP_3$ production by PLCdelta |
| kappa_delta | 1.5 µM | Inhibition constant of PLCdelta by $IP_3$ |
| K_delta | 0.1 µM | $Ca^{2+}$ affinity of PLCdelta |
| Omega_5P | 0.05 /s | Maximal rate of $IP_3$ degradation by IP-5P |
| K_D | 0.7 µM | $Ca^{2+}$ affinity of $IP_3$-3K |
| K_3K | 1.0 µM | $IP_3$ affinity of $IP_3$-3K |
| O_3K | 4.5 µM/s | Maximal rate of $IP_3$ degradation by $IP_3$-3K |
| F | 0.09 µM/s | GJC $IP_3$ permeability |
| I_Theta | 0.3 µM | Threshold gradient for $IP_3$ diffusion |
| omega_I | 0.05 µM | Scaling factor of diffusion |
| Omega_A | 0.6 /s | Gliotransmitter recycling rate |
| Omega_e | 60 /s | Gliotransmitter clearance rate |
| alpha | 0.0 | Gliotransmission nature |
| duration | 4 s | Total simulation time |

**Table S2**: Simulation parameters used in the neural network model.
